# VirB, a key transcriptional regulator of *Shigella* virulence, requires a CTP ligand for its regulatory activities

**DOI:** 10.1101/2023.05.16.541010

**Authors:** Taylor M. Gerson, Audrey M. Ott, Monika MA. Karney, Jillian N. Socea, Daren R. Ginete, Lakshminarayan M. Iyer, L. Aravind, Ronald K. Gary, Helen J. Wing

## Abstract

The VirB protein, encoded by the large virulence plasmid of *Shigella* spp., is a key transcriptional regulator of virulence genes. Without a functional *virB* gene, *Shigella* cells are avirulent. On the virulence plasmid, VirB functions to offset transcriptional silencing mediated by the nucleoid structuring protein, H-NS, which binds and sequesters AT-rich DNA, making it inaccessible for gene expression. Thus, gaining a mechanistic understanding of how VirB counters H-NS-mediated silencing is of considerable interest. VirB is unusual in that it does not resemble classic transcription factors. Instead, its closest relatives are found in the ParB superfamily, where the best-characterized members function in faithful DNA segregation before cell division. Here, we show that VirB is a fast-evolving member of this superfamily and report for the first time that the VirB protein binds a highly unusual ligand, CTP. VirB binds this nucleoside triphosphate preferentially and with specificity. Based on alignments with the best-characterized members of the ParB family, we identify amino acids of VirB likely to bind CTP. Substitutions in these residues disrupt several well-documented activities of VirB, including its anti-silencing activity at a VirB-dependent promoter, its role in generating a Congo red positive phenotype in *Shigella*, and the ability of the VirB protein to form foci in the bacterial cytoplasm when fused to GFP. Thus, this work is the first to show that VirB is a bona fide CTP-binding protein and links *Shigella* virulence phenotypes to the nucleoside triphosphate, CTP.

**Importance:** *Shigella* species cause bacillary dysentery (shigellosis), the second leading cause of diarrheal deaths worldwide. With growing antibiotic resistance, there is a pressing need to identify novel molecular drug targets. *Shigella* virulence phenotypes are controlled by the transcriptional regulator, VirB. We show that VirB belongs to a fast-evolving, primarily plasmid-borne clade of the ParB superfamily, which has diverged from versions that have a distinct cellular role – DNA partitioning. We are the first to report that, like classic members of the ParB family, VirB binds a highly unusual ligand, CTP. Mutants predicted to be defective in CTP binding are compromised in a variety of virulence attributes controlled by VirB. This study i) reveals that VirB binds CTP, ii) provides a link between VirB-CTP interactions and *Shigella* virulence phenotypes, and iii) broadens our understanding of the ParB superfamily, a group of bacterial proteins that play critical roles in many different bacteria.

## Introduction

In *Shigella flexneri,* an intracellular bacterial pathogen and the causative agent of bacillary dysentery (1–4), the transcriptional regulator VirB is produced upon entry into the host in response to human body temperature (5). VirB directly or indirectly leads to the upregulation of about 50 genes located on the large virulence plasmid, pINV (6–8), including those encoding the type III secretion system, other crucial virulence-associated factors (OspZ, OspD1, and IcsP (9–11), and the transcriptional activator, MxiE, and its co-activator, IpgC (12). Consequently, VirB-dependent gene regulation is essential for *Shigella* virulence (7). In the late 1970s/early 1980s, *Shigella* virulence was found to directly correlate with the ability of colonies to bind the organic dye Congo red (13–15). Mutants lacking *virB*, exhibit a Congo red negative (CR-) phenotype (16, 17), providing an easy way to evaluate the activity of VirB and assess *Shigella* virulence.

VirB does not function like a traditional transcription factor, rather, it functions as an anti-silencing protein, alleviating transcriptional silencing mediated by the histone-like nucleoid structuring protein, H-NS (9, 11, 18–23). H-NS coats and condenses DNA by binding to the minor groove of AT-rich DNA, which is a common feature of horizontally acquired DNA (24, 25). Thus, H-NS serves as a xenogeneic silencer of newly acquired DNA (26–30), which includes key virulence genes in many important bacterial pathogens (31–35). Alleviation of H-NS-mediated silencing is caused by VirB engaging its DNA recognition site and spreading along DNA (22), which is thought to remodel H-NS-DNA complexes, allowing previously inaccessible or transcriptionally nonpermissive DNA to be bound by RNA polymerase (23). Recent work suggests that anti-silencing or the remodeling of H-NS-DNA complexes is likely mediated by a localized loss of negative DNA supercoils, which is triggered when VirB engages its site and spreads along DNA, causing a stiffening of the helix (36).

VirB is not related to classic bacterial transcriptional regulators but is a member of the ParB superfamily (18, 37–39). Most members of the ParB superfamily are characterized to play key roles in chromosome or plasmid segregation prior to cell division (40–42). While VirB does not function in DNA segregation (39, 43), we hypothesize that its evolutionary relationship to ParB proteins will provide insight into its mechanism of anti-silencing. ParB proteins load onto DNA at palindromic DNA recognition sites, called *parS* (44–46), and condense adjacent DNA regions into large nucleoprotein complexes (47). Evidence suggests that these nucleoprotein complexes are formed by the lateral spreading of ParB proteins along DNA into regions flanked by *parS* (48–51). After ParB proteins establish partitioning complexes, an ATPase, ParA, is recruited to the complex and interacts with the N-terminus of ParB to move sister replicons to the poles of the cell through a ratchet-like mechanism (52–54). Of note, functional GFP-ParB fusions form fluorescent foci at the pole prior to cell division. Recently, some members of the ParB superfamily have been shown to bind and hydrolyze CTP. The CTP ligand is needed for an open-to-closed conformational change, which allows ParB to dissociate from the *parS* site and spread along DNA (55–57).

Like ParB proteins, VirB recognizes a palindromic *parS*-like DNA binding site (18, 20, 22, 39, 58), and multimers of VirB form on DNA both *in vitro* and *in vivo* (37, 38, 59). Both activities are essential for the transcriptional anti-silencing of virulence genes by VirB (22, 37–39, 58, 59). However, there are clear differences between VirB and ParB proteins too. Aside from different cellular functions (11, 20, 60), the *virB* locus lacks a *parA*-like gene (43) and the VirB protein does not require a ParA-like partner protein in order to function (18, 43, 61). The recent discovery that some ParB members function as CTP-binding proteins raises questions as to whether VirB binds CTP or some other NTP and what effect this may have on transcriptional anti-silencing. Thus, the overarching goals of this study were to i) gain further insight into the ParB superfamily and the relationship between classic ParB proteins and VirB, ii) determine if VirB is capable of binding CTP, and if so, iii) characterize how CTP binding may impact the virulence phenotypes controlled by VirB.

## Results

### VirB is a member of a fast-evolving clade within the ParB superfamily

To understand the provenance and potential biochemical functions of *Shigella* VirB, we recovered both immediate and more distant homologs (see Materials and Methods). Searches typically retrieved several VirB-related ParB superfamily members, e.g., *Myxococcus xanthus* PadC and *Agrobacterium vitis* ParB from the pAtS4c plasmid as top hits, and then recovered the primary genome-encoded classic chromosome partitioning ParB proteins. A representative sequence set was used to construct a multiple alignment to analyze the sequence conservation patterns and for phylogenetic analysis. The phylogenetic trees revealed a consistent and well-supported topology in which *Shigella* VirB and a group of predominantly proteobacterial proteins formed a distinct clade to the exclusion of the remaining classic bacterial ParB proteins (Figure 1A & S1). This separation was corroborated by a clear bimodal distribution of the scaled and centered distances of the classic ParB from cellular genomes and VirB-related proteins. The former constituted a peak at the left of the distribution with short distances, and the latter right peak corresponded to large distances (Figure 1B). The mean scaled and centered distances are significantly higher for the VirB-like clade as opposed to the classic cellular-genome-encoded ParBs (p< 2.2 x 10^-16^), indicating that the VirB-like clade is fast-evolving relative to the classic ParBs (the slow-evolving clade).

**Figure 1.**
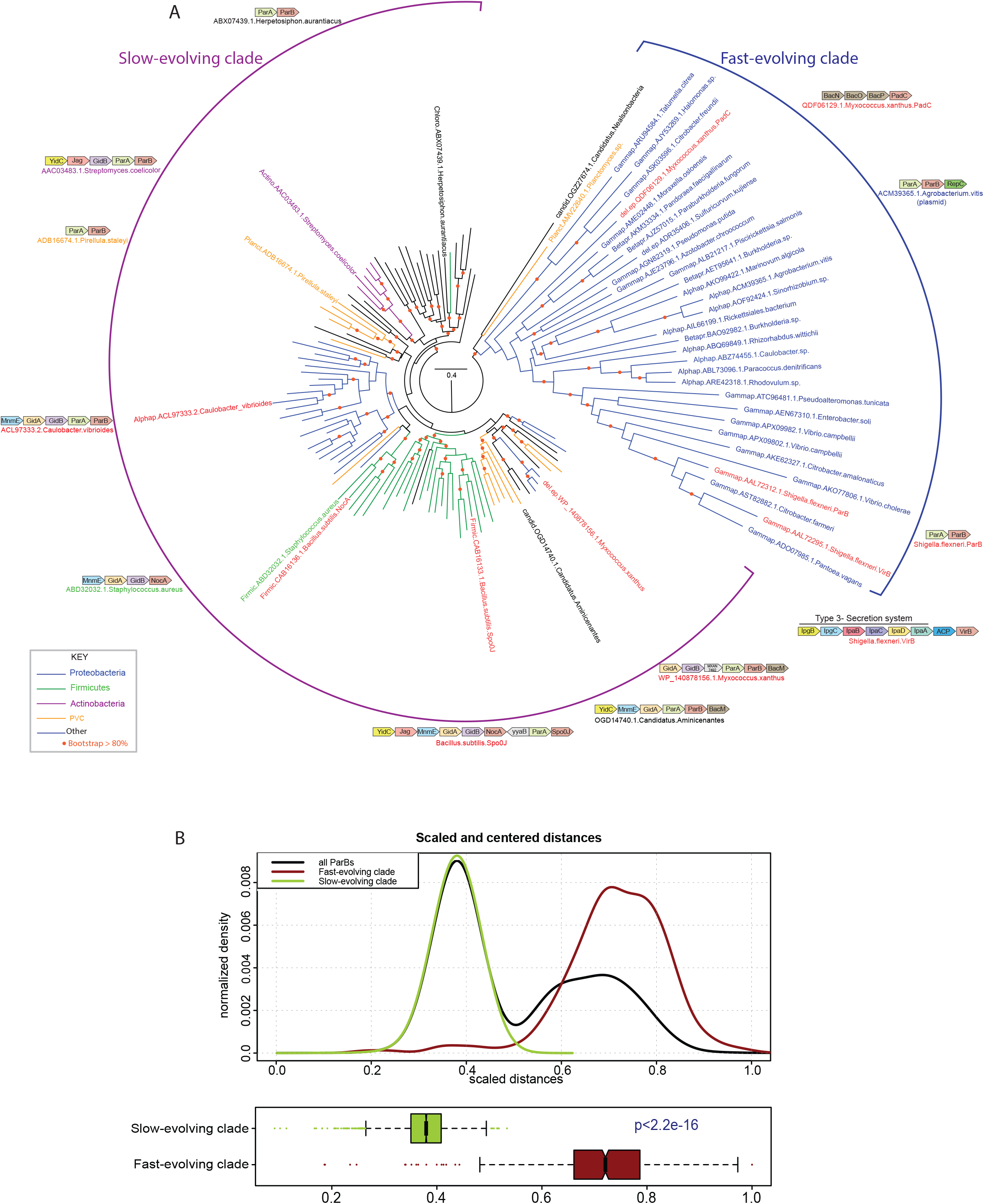
Maximum likelihood tree of ParB protein sequences including the VirB-like fast-evolving sequences. (A) Maximum likelihood tree of ParB protein sequences including the VirB-like fast-evolving sequences. The tree was computed using IQtree and bootstrapped with 1000 replicates. Nodes and branches corresponding to widely represented phylogenetic clades are colored according to the key. Protein sequences are denoted with their clade, gene and species name. The slow- and fast-evolving clades are marked and representative gene neighborhoods of distinct clades of the tree are shown. Genes in gene neighborhoods are shown as box arrows with the arrowhead pointing to the gene at the 3’ end. Clade abbreviations include: Actino: Actinomycetes, Alphap: Alpha-proteobacteria, Betapr: Beta-proteobacteria, candid: Dark-matter bacteria, Chloro: Chloroflexi, del.ep: Delta/epsilon proteobacteria, Firmic: Firmicutes, Gammap: Gamma-proteobacteria, Planct: Planctomycetes. (B) Differential evolutionary rates derived from pairwise centered and scaled maximum-likelihood distances shown as a kernel density and box plots.

These cladal differences are also reflected in gene-neighborhood contexts (Figure 1A & S1). Across bacteria, classic ParBs from cellular genomes tend to be encoded in largely conserved gene-neighborhoods along with genes encoding: (i) GidA and MnmE, both of which are involved in the synthesis of the carboxymethylaminomethyl modification of uridine 34 of tRNAs (62, 63); (ii) GidB, the 16S rRNA guanidine 535 N7 methylase (64, 65); (iii) YidC, the membrane protein assembly factor (66, 67); (iv) Jag, an RNA-binding protein with KH and R3H domains that regulates the expression of the cell-division protein FtsA post-transcriptionally (68); (v) ParA, the ATPase partner of ParB required for chromosome segregation (46, 53, 69, 70). Less frequently, these neighborhoods might also code for one or more genes for the β-helix cytoskeletal protein bactofilin, implicated in cell division along with ParA and ParB (71–74). In sharp contrast, other than the neighborhood association with genes encoding ParA and, on rare occasions, bactofilin (e.g., *M. xanthus* PadC), the fast-evolving VirB-like clade lacks the remaining associations of classic ParBs. Additionally, the fast-evolving clade is primarily found on free plasmids or plasmids integrated into cellular genomes. These findings suggest that, with the exception of partnering with ParA, the fast-evolving VirB-like ParBs do not have the genetic and physical interaction constraints that act on the classic ParBs, which has allowed them to evolve more rapidly.

Both the classic ParBs and the VirB-like clade possess a conserved domain architecture with 4 evolutionarily distinct domains including (i) a ParB NTP-binding/hydrolyzing domain; (ii) a DNA-binding helix-turn-helix domain; (iii) a tetra-helical bundle comprised of two α-α-hairpins; (iv) a C-terminal domain likely playing a role both in dimerization and DNA binding. Together they form a dimer with a central aperture through which DNA can be threaded. Additionally, they contain an unstructured N-terminal tail with two conserved arginines that are proposed to act as “arginine fingers” to facilitate ATP-hydrolysis by the ParA partner protein. This N-terminal tail is a diagnostic feature of ParB superfamily proteins that interacts with a ParA partner (Figure S2). Notably, we found that this tail has diverged considerably and lost the arginines in VirB, consistent with the lack of a ParA partner interaction (Figure S2). However, VirB is nested in the tree with versions that do interact with a ParA protein (e.g., ParB from *Shigella flexneri* plasmid pCP301). This suggests a two-step model for the emergence of VirB from a ParB-like precursor. First, release from the constraints typical of the genomic classic ParBs allowed the plasmid-borne versions to rapidly evolve and diverge. This divergence allowed the exploration of new functions, such as transcriptional regulation, which utilized the ancestral DNA-binding properties of ParB for a different role. This favored the loss of the determinants of its original function, i.e., the ParA-interaction residues. Indeed, multimer modeling via the AlphaFold2 program indicated that while the slow-evolving ParBs form a predicted complex with ParA via the N-terminal tail, *Shigella* VirB did not yield any such complex.

### The *Shigella* anti-silencing protein VirB binds a novel ligand, CTP

Despite its overall rapid divergence, all members of the fast-evolving clade, including VirB, retain key NTP-binding determinants conserved across the ParB superfamily (Figure S2) and at least one member of this clade, *M. xanthus* PadC, binds CTP like the classic ParBs (55–57). Hence, we next investigated if VirB binds CTP. Using isothermal titration calorimetry (ITC), CTPγS, CTP, or UTP was titrated into a cell containing purified VirB protein, and differential power (DP) was plotted against time (Figure 2).

**Figure 2.**
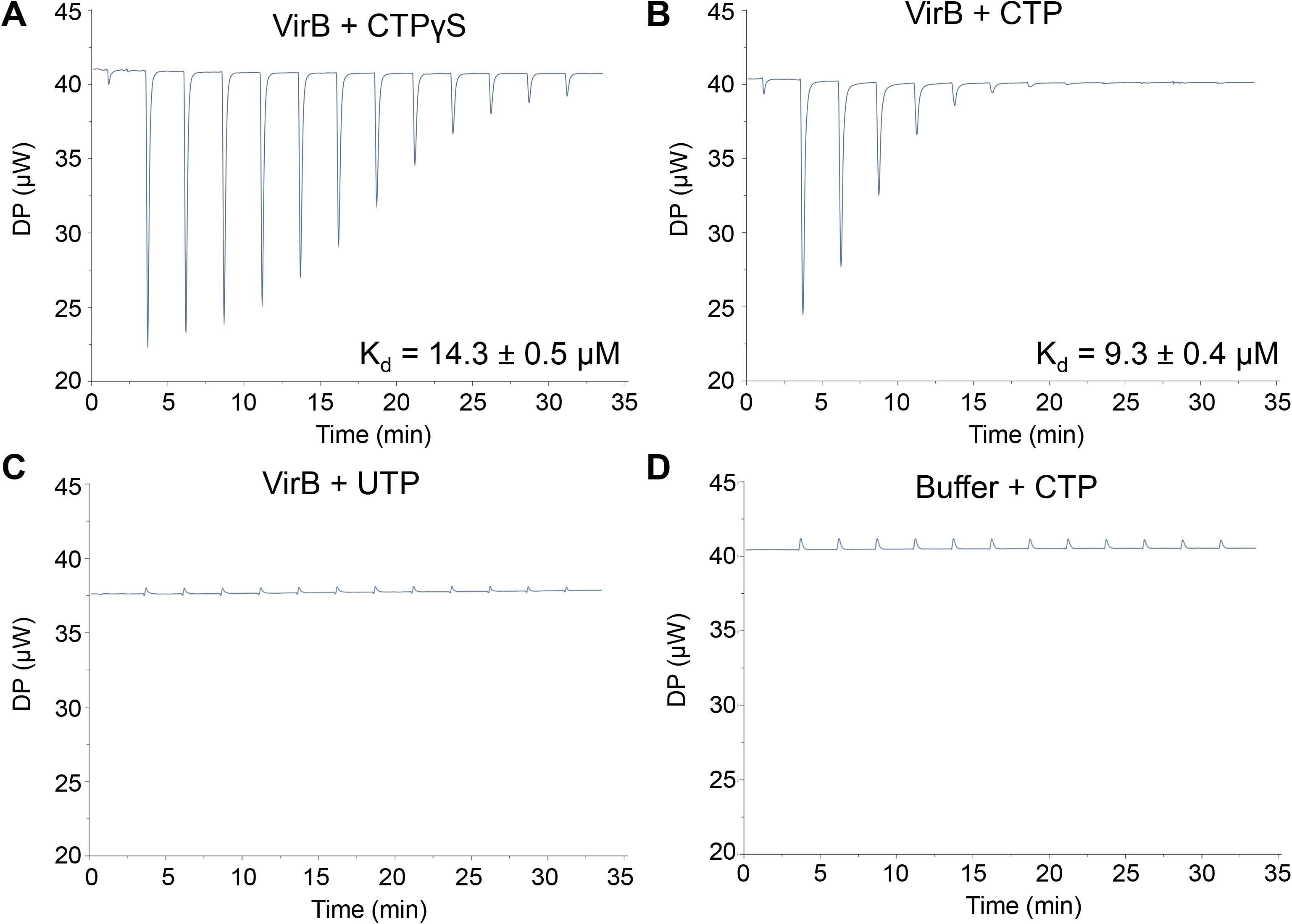
The *Shigella* anti-silencing protein VirB binds a novel ligand, CTP. ITC measurements with (A) 90 µM VirB-His6 and 3 mM CTPγS, (B) 45 µM VirB-His6 and 3 mM CTP, (C) 45 µM VirB-His6 and 3 mM UTP, and (D) VirB binding buffer and 3 mM CTP. In panels B and C, the VirB protein concentration was lowered to 45 µM to reach saturation and conserve protein.

During initial experiments, non-hydrolyzable CTP, CTPγS, was titrated into the VirB-containing cell to eliminate possible interference from any potential nucleotide hydrolysis. The addition of CTPγS produced exothermic binding events (downward-facing peaks) that gradually diminished as binding sites became saturated (Figure 2A). The data were fit to a "one set of sites" (i.e., non-cooperative) binding model, and VirB was calculated to bind CTPγS with a Kd value of 14.3 µM ± 0.5 µM, values similar to those calculated for Spo0J (*B. subtilis*), a classic ParB member (10.3 ± 1.2 µM) (56). Because VirB was not fully saturated by CTPγS after 13 injections, we decreased the VirB concentration by two-fold in subsequent experiments. Commercially available CTP was found to bind VirB with affinities similar to those seen with CTPγS (Kd = 9.3 µM ± 0.4 µM; Figure 2B). To determine if VirB binds other pyrimidine triphosphates available in the cell, UTP was used. In contrast to CTP, UTP showed no binding signatures (Figure 2C), resembling the buffer-only controls (Figure 2D & Figure S3A & B). Failure of VirB to bind UTP strongly supports the conclusion that VirB binds CTP with specificity. After the binding sites on VirB were saturated, subsequent injections of CTP produced no thermal signal (Figure 2B), suggesting VirB is incapable of hydrolyzing CTP under these experimental conditions. Thus, VirB, the anti-silencing protein of *S. flexneri* virulence genes, is a bona fide CTP-binding protein and binds CTP with affinities similar to those exhibited by classic ParBs (55–57).

### VirB preferentially binds CTP over other NTP ligands found in the cell

Our ITC experiments revealed that VirB binds CTP, but not UTP (Figure 2). However, to conclusively test whether VirB binds any other nucleoside triphosphates available in the cell, we exploited the differential radial capillary action of ligand assay (DRaCALA) with cold competitor NTPs (75). Briefly, DRaCALA is based on the premise that a small, radiolabeled ligand will diffuse on nitrocellulose, but, if bound to a protein of interest, the protein:ligand complex will not diffuse and remain at the original point of contact, resulting in a tight spot of radioactive signal (dark inner core). In our study, we incubated purified VirB protein with radiolabeled (hot) CTP and, where appropriate, excess unlabeled (cold) NTP ligands (CTP, UTP, ATP, or GTP). Reaction mixtures were then spotted onto dry nitrocellulose membranes and air-dried, prior to imaging.

As expected, in the absence of VirB, but in the presence of radiolabeled (hot) CTP, no dark inner core was detected, yet in the presence of both, a dark inner core was seen. These data indicate that VirB binds the radiolabeled CTP (Figure 3A), supporting our ITC data. Next, cold competitor NTPs were included in the reaction mix at a 100,000-fold molar excess. When excess cold CTP was added to the reaction mix, the radioactive signal was diffuse, indicating that the cold competitor CTP had competed for VirB binding (Figure 3A). In support of this, a significant decrease in the fraction of radiolabeled CTP bound was quantified, relative to that found in the absence of the competitor (Figure 3B). In contrast, when excess cold UTP, ATP, or GTP was added to the reaction mixture, no competition for the VirB protein was observed (Figure 3A & B). These results conclusively demonstrate that VirB binds CTP preferentially and with specificity.

**Figure 3.**
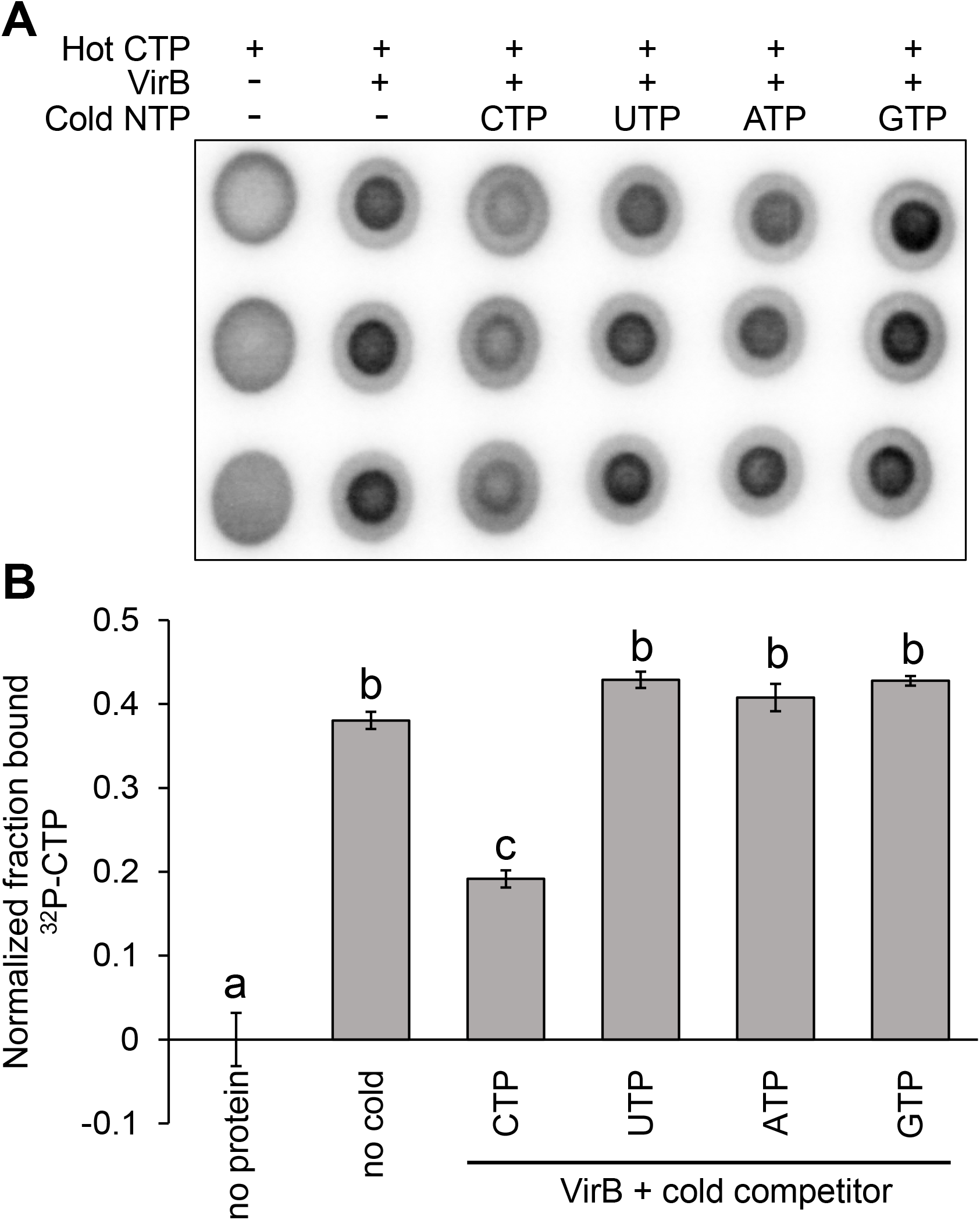
VirB preferentially binds CTP over other NTP ligands found in the cell. (A) DRaCALA images of competition assays assessing the ability of 500 µM indicated cold NTP to compete with binding interactions between 5 nM ^32^P-CTP and 50 µM VirB-His_6_. Ligand binding is indicated by a darker inner core staining due to rapid immobilization of the VirB-CTP complex. (B) Graph of normalized fraction bound (F_B_) calculated for each sample in Figure 3A. Assays were completed with three technical replicates and three biological replicates. Representative data are shown. Significance was calculated using a one-way ANOVA with post hoc Bonferroni, *p* < 0.05. Lowercase letters indicate statistical groups. Complete statistical analysis provided in Supplementary Table S3.

### Identification of key VirB residues predicted to be required for CTP binding

With the discovery that VirB binds CTP, we next wanted to identify amino acid residues in VirB predicted to be required for CTP binding. A multiple sequence alignment (MSA) that included VirB and the well-characterized ParB proteins demonstrated to bind CTP (*B. subtilis* Spo0J (56), *C. crescentus* ParB (57), and *M. xanthus* PadC (55)) allowed us to compare residues previously implicated in CTP binding (Figure 4A & S2). These are located in two regions (described as Box I & Box II; (55)) near the N-terminus of their respective proteins (55–57). Residues in the motif ELXXSIXXXGXXXP (Box I) were characterized to interact with the cytosine base of CTP and thus, are critical for CTP binding specificity (55, 76). Whereas residues in the motif GERRXRA (Box II) were characterized to interact with triphosphate motifs of CTP, playing a role in stabilizing this interaction (55, 76). Our MSA (Figure 4A & S1) showed that in Box I, two residues (I65 & F74) correspond to similar residues found in the classic ParBs (Figure 4A). Additionally, T68, located in Box I, corresponds to a serine in the slow-evolving ParBs, which hydrogen bonds to the cytosine base (Figure 4A), suggesting T86 might play a comparable role in VirB. In Box II, three residues (G91, R93, & R94) were identified that were fully conserved, when compared to classic ParBs (Figure 4A). Thus, six residues of interest I65, T68, F74, G91, R93, & R94 were identified. These were explored further in this study.

**Figure 4.**
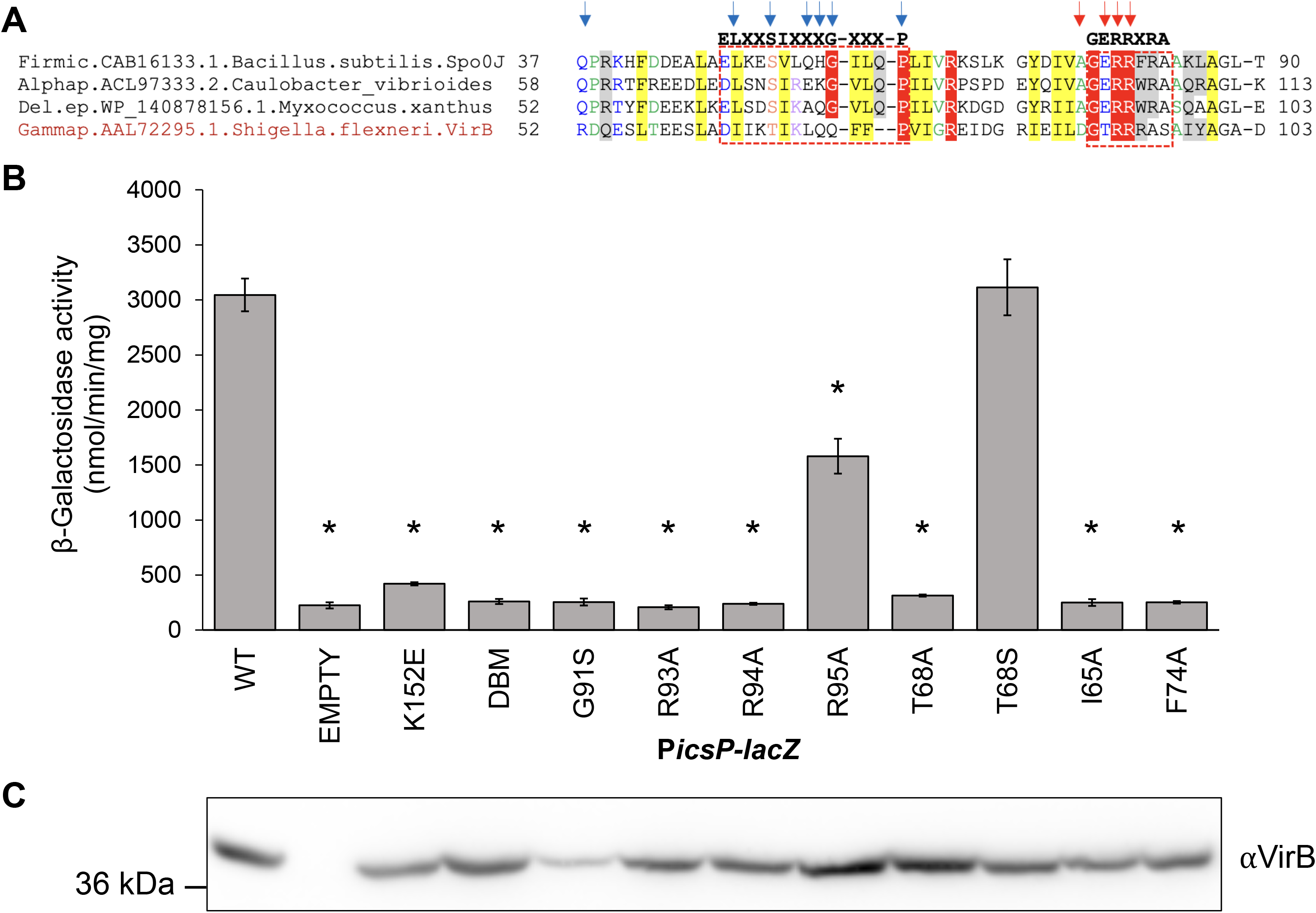
Key residues in the predicted CTP binding pocket are required for the anti-silencing activity of VirB. (A) Multiple sequence alignment (MSA) of classic ParBs that bind CTP and VirB, generated using the Mafft program with a local pair algorithm. Full alignment and additional details provided in Figure S2. Taxonomic clade name, Genbank accession, and species name separated by dots are denoted. Consensus abbreviations and coloring scheme are as follows: hydrophobic (FWYILVACM), aromatic (FWY) and aliphatic (ILV) residues shaded yellow; charged (DEHKR) and basic (KRH) residues colored magenta, big (LIFMWYERKQ) residues shaded grey, alcohol-group (ST) residues colored red, polar (STECDRKHNQ) residues colored blue, small (AGSCDNPTV) residues colored green, and tiny (GAS) residues shaded green. Fully conserved residues are shaded red. Arrows indicate nucleotide-binding residues (blue) and catalytic residues (red). (B) β-Galactosidase assay used to assess the regulatory activity of pBAD-VirB derivatives at the VirB-dependent *icsP* promoter. Significance was calculated using a one-way ANOVA with post hoc Tukey HSD, *p* < 0.05. *, statistically significant compared to wild-type. Complete statistical analysis is provided in Supplementary Table S4. (C) Western Blot analysis using an anti-VirB antibody to assess protein production of pBAD-VirB mutants alongside a SeeBlue^TM^ Plus2 Prestained Standard. Assays were completed with three biological replicates and repeated three times. Representative data are shown.

### Key residues in the predicted CTP binding pocket are required for the anti-silencing activity of VirB

Key VirB residues predicted to bind CTP were targeted for substitution. A suite of *virB* mutant alleles was generated and introduced into the inducible pBAD expression plasmid to generate the following VirB mutants; I65A, T68A, F74A, G91S, R93A, and R94A. Additionally, VirB T68S and VirB R95A were created because we reasoned that T68S would retain the potential for hydrogen bonding at this position, whereas R95A substitutes a non-conserved residue that lies adjacent to a patch of fully conserved core residues.

To assess the impact of these substitutions on VirB-dependent gene regulation, we measured the ability of these VirB mutants to regulate a well-characterized VirB-regulated promoter, P*icsP*. Inducible plasmids each carrying a pBAD-*virB* mutant were introduced into a *virB* mutant derivative of *Shigella flexneri* (AWY3) carrying the P*icsP-lacZ* transcriptional reporter (9, 22), and β-galactosidase activity was measured in cultures following induction. As expected, wild-type VirB had high levels of β-galactosidase activity, indicating that VirB was able to regulate P*icsP*, whereas the empty control and two well-characterized DNA binding mutants (a single and double mutant; (11, 20)) each exhibited low levels of β-galactosidase activity (Figure 4B). Strikingly, the following VirB mutants I65A, T68A, F74A, G91S, R93A, and R94A exhibited a significant decrease in β-galactosidase activity when compared to wild-type and displayed activities similar to the negative controls (Figure 4B). However, VirB T68S showed no significant decrease (Figure 4B). The loss of activity by VirB T68A, but the retention by VirB T68S, suggests that the anti-silencing activities of VirB require residue 68, which is predicted to bind CTP, to participate in hydrogen bond formation. Interestingly, VirB R95A showed significantly lower levels of β-galactosidase activity when compared to wild-type VirB, although this activity was significantly higher than that displayed by most of the VirB mutants (Table S3). Thus, even though R95 is not a residue that is conserved in other family members, VirB R95A displayed intermediate levels of anti-silencing activity in our assays (Figure 4B).

Importantly, western analyses of cell pellets generated from cultures grown identically to those used in our β-galactosidase assays, revealed that all proteins, with one exception, VirB G91S, were made at wild-type levels or higher under assay conditions. Since the instability of VirB G91S raised concerns about whether this derivative was structurally sound, it was eliminated from subsequent studies. Taken together, our P*icsP* promoter activity assays and western analyses show that key residues located in the predicted CTP binding pocket of VirB, are required for its transcriptional anti-silencing activity.

### Key residues in the predicted CTP binding pocket are required for *Shigella* virulence phenotype

To further examine the effect that our VirB mutants have on virulence phenotypes, we next assessed the Congo red binding of *Shigella* cells expressing the *virB* mutant derivatives. The ability of *Shigella* colonies to bind Congo red is positively correlated with their virulence properties (13, 14). Congo red binding assays were performed using a *virB* mutant derivative of *Shigella flexneri* (AWY3) expressing each of the inducible *virB* mutant alleles from pBAD-*virB* derivatives, described previously.

Briefly, Congo red phenotypes (CR^+^ = red = virulent, and CR^-^ = white = avirulent) for each mutant were observed on Congo red plates under either non-inducing or inducing conditions (0.2% D-glucose or L-arabinose, respectively). Under non-inducing conditions, all colonies displayed a CR^-^ phenotype (Figure S4). As expected, under inducing conditions, wild-type VirB appeared CR^+^ phenotype (Figure 5A). In contrast, mutants found to be incapable of regulating the *icsP* promoter (I65A, T68A, F74A, R93A, & R94A), displayed a CR^-^ phenotype, similar to that seen under non-inducing conditions (Figure S3 and quantified in Figure 5A & B). These findings are consistent with the idea that native residues at these positions are needed for *Shigella* virulence phenotypes. Notably, in these assays, VirB T68S and R95A exhibited CR^+^ phenotypes similar to wild-type (Figure 5A & B), whereas the annotated DNA-binding mutant K152E displayed an intermediate level of Congo red binding, statistically different from both the wild-type and those displaying a true CR^-^ phenotype. Collectively, these results show that key residues in VirB predicted to bind CTP are required for Congo red binding, a phenotype positively correlated with *Shigella* virulence.

**Figure 5.**
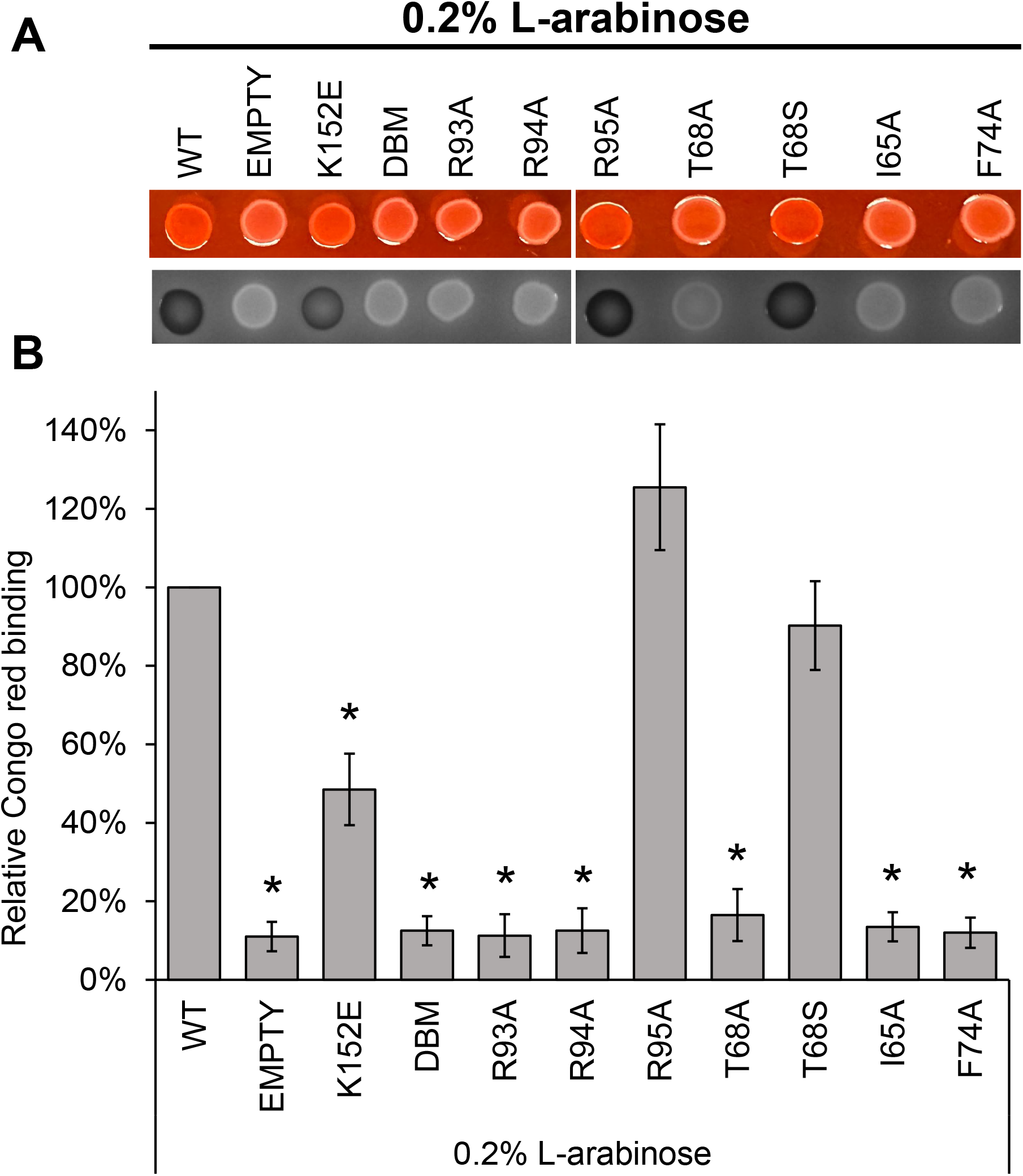
Key residues in the predicted CTP binding pocket are required for *Shigella* virulence phenotype. (A) Congo red binding by *S. flexneri virB*::Tn5 harboring pBAD-VirB derivatives under inducing conditions. Images were captured using visible light (top) and blue light (Cy2) (bottom). (B) Quantitative analysis of Congo red binding *S. flexneri virB*::Tn5 harboring pBAD-VirB derivatives (induced). Relative Congo red binding was calculated as [(OD_498_/OD_600_)/(average (OD_498_/OD_600_)_2457T pBAD_)] × 100. Assays were completed with three biological replicates and repeated three times. Representative data are shown. Significance was calculated using a one-way ANOVA with post hoc Tukey HSD, *p* < 0.05. *, statistically significant compared to wild-type. Complete statistical analysis is provided in Supplementary Table S5.

### Key residues in the predicted CTP binding pocket are required VirB focus formation *in vivo*

Members of the ParB superfamily, to which VirB belongs, show a discrete subcellular localization pattern within the bacterial cell during faithful plasmid/chromosome segregation, an activity associated with engaging its DNA recognition site (46, 47, 77–80). Even though VirB has an entirely different cellular role, a recent study in our lab discovered that VirB also forms discrete foci within *Shigella* cells, which are dependent upon VirB binding to its recognition site (81). We next investigated what effect VirB mutants would have on discrete focus formation when fused to superfolder green fluorescent protein (sfGFP) (82, 83). To test this, the 5’ end of five *virB* mutant alleles encoding substitutions in either T68 (both Ala and Ser), R93, R94, or R95, were introduced into the pBAD promoter located on the low-copy (∼15-20 copies per cell) plasmid, pGB682. The resulting plasmids were induced in a *virB* mutant derivative of *Shigella flexneri* (AWY3) (81). The distribution of foci formed was quantified using the number of maxima detected per cell. Based on our previous study (81), any observed foci would likely be caused by the interaction of VirB with the large virulence plasmid.

As expected, in cells producing GFP only, there was a diffuse GFP signal, with 98% of cells having a single focus (maxima), whereas the empty plasmid control gave no GFP signal (Figure 6A & B). Cells producing GFP-VirB displayed discrete focus formation, with a majority of cells having 2-3 foci per cell, as described previously (81). This effect is likely dependent upon VirB engaging its DNA recognition site (Figure 6A & B), supported by a fusion carrying two substitutions in the DNA-binding domain (GFP-VirB DBM) exhibiting a diffuse signal like the GFP control (Figure 6A & B). When GFP was fused to VirB with alanine substitutions at key residues in the putative CTP binding pocket (R93A, R94A, or T68A) no foci were observed, also resembling the GFP control. In contrast, GFP-VirB R95A and T68S displayed discrete focus formation, with most cells forming two foci, similar to the wild-type fusion (Figure 6A & B). This is consistent with R95 being non-conserved but proximal to the CTP binding pocket residues and T68S potentially restoring hydrogen bonding at this position. In sum, these findings demonstrate that R93, R94, and T68, residues in the CTP binding pocket, are required for VirB focus formation *in vivo.* This raises the possibility that GFP-VirB R93A, R94A, and T68A are defective in DNA binding *in vivo*, although further work is needed to test this conclusively.

**Figure 6.**
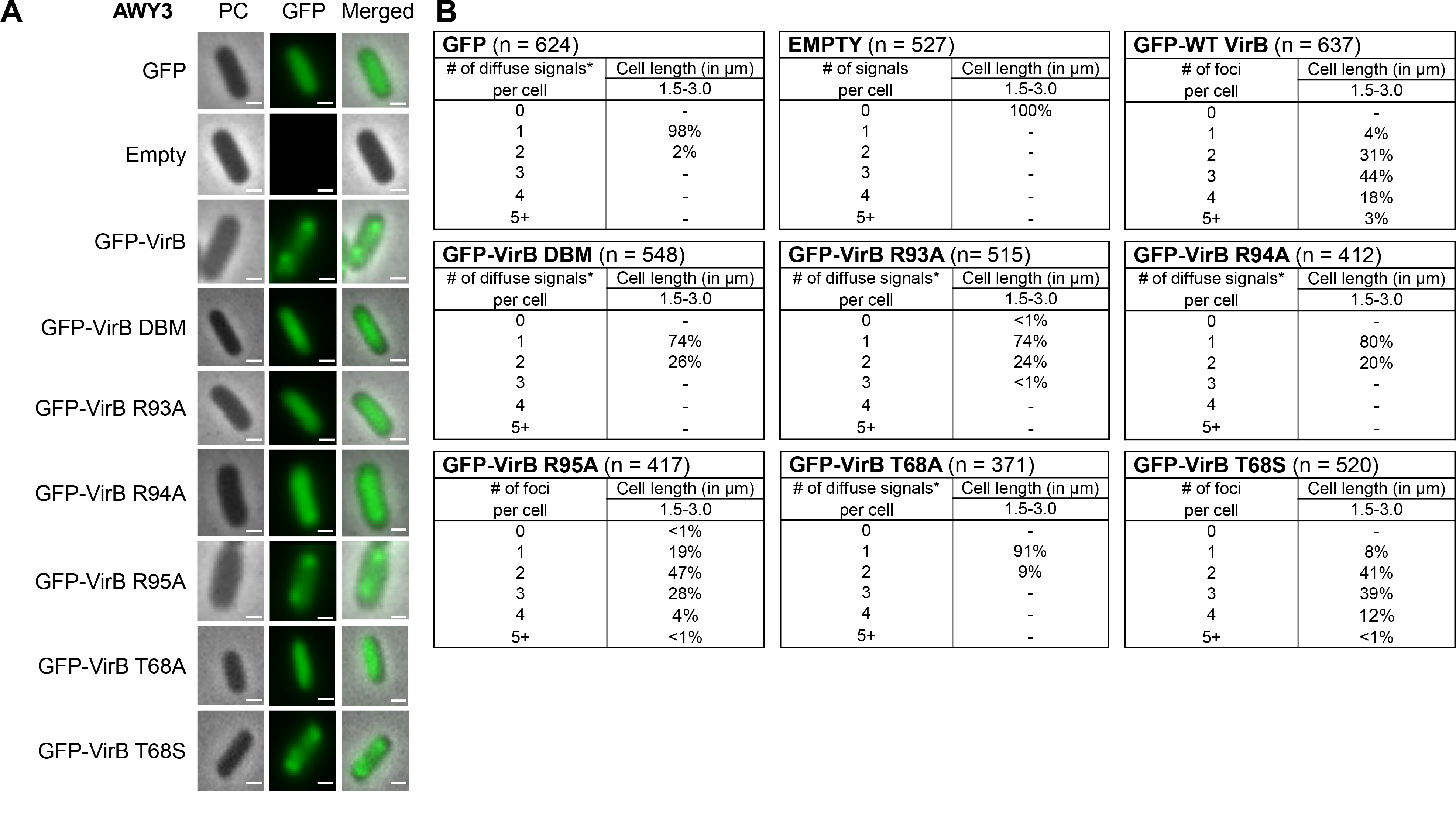
Key residues in the CTP binding pocket are required for VirB focus formation *in vivo*. (A) Live-cell imaging of GFP-VirB derivatives in a *virB* mutant strain of *S. flexneri* (AWY3) induced with 0.02% L-arabinose. Phase-contrast, PC, (left column), fluorescence (middle column) (GFP row, 28-ms exposure; GFP-VirB derivatives and empty rows, 98-ms exposure), and merged (right column). Bars represent 1 µM. (B) Quantification of fluorescent signals observed during live-cell imaging of GFP-VirB derivatives using MicrobeJ. Within tables, a hyphen indicates that no cells were found in this category in any of the images captured; *, maxima detected by MicrobeJ. Complete statistical analysis is provided in Supplementary Table S6 and field-of-view images are found in Figure S5.

## Discussion

In this work, we focus on the transcriptional anti-silencing protein, VirB, which functions to offset H-NS-mediated silencing of *Shigella* virulence genes on the large virulence plasmid. We show that VirB is a member of a fast-evolving clade of ParB proteins that is distinct from the slow-evolving versions that function in the segregation of the primary cellular chromosome (Figure 1). Using two different approaches (ITC and DRaCALA), we show that, like members of the slow-evolving clade, VirB binds a highly unusual ligand, CTP, preferentially and with specificity (Figure 2 & 3). VirB residues likely required for CTP binding were identified using amino acid alignments to the best-characterized members of the ParB family and were shown to be necessary for (i) VirB-dependent anti-silencing of virulence genes, (ii) generation of a Congo red positive phenotype, which correlates with *Shigella* virulence, and (iii) the ability of the VirB protein to form discrete foci when fused to GFP (Figure 4, 5, & 6). Consequently, our work provides a new perspective on the ParB superfamily and is the first to link virulence attributes of *Shigella* to the highly unusual nucleoside triphosphate ligand, CTP. Our work also raises questions about the role that CTP plays in the VirB-dependent mechanism of virulence gene regulation.

To the best of our knowledge, the phylogenetic analysis of the ParB superfamily, included in this work, is the first to include VirB. Our analysis reveals that this superfamily is comprised of two distinct clades, one that contains the classic ParBs and a second, fast-evolving clade, containing *Shigella* VirB (Figure 1A). The latter clade is primarily comprised of plasmid-encoded proteins found in Proteobacteria (Figure 1B). Our studies highlight that at least two members of this fast-evolving clade have become separated, in terms of their gene-neighborhoods, from *parA* (which encodes the ATPase that functions in faithful DNA segregation), i.e., *M. xanthus* PadC [*padC* is the fourth gene in an operon encoding bactofilins; (84)] & *S. flexneri* VirB [*virB* is monocistronic and located immediately after the *ipa* operon, which encodes type III effector proteins, on pINV; (6, 7)] (Figure 1 & S1). While PadC of *M. xanthus* still interacts with a ParA protein, encoded by the *parA* linked to *M. xanthus parB* (84), our analysis reveals that VirB has lost the N-terminal arginines which mediate interactions between ParB and ParA in other members of the family (Figure S2). For VirB, this loss of functional connection with a ParA-like partner protein has gone hand-in-hand with the acquisition of a new function (43, 85). We propose that the rapid evolution of the plasmid-borne clade allowed these proteins to adopt new functions based on their existing biochemistry, i.e., NTP and DNA binding. The latter property was eminently suited for VirB, at least, to adopt a new role as a transcription factor.

A major thrust of our research is to understand the molecular mechanism that is used by VirB to control *Shigella* virulence (11, 20, 22, 36, 58, 81). In pursuit of this, we have drawn from work on the ParB proteins and how they interface with DNA (48, 51, 69, 78, 86, 87). VirB and ParB proteins engage DNA as homodimeric proteins, through the recognition of a similar inverted repeat (18, 38, 39, 44, 88). Recently, though, members of ParB family (*B. subtilis* Spo0J (56), *C. crescentus* ParB (57), *M. xanthus* ParB (55)) have been identified to bind and hydrolyze a unique, nucleoside triphosphate, CTP, in order to switch between open and closed clamp conformations while spreading along DNA (55–57). Here, we are the first to report that *Shigella* VirB binds CTP preferentially and with specificity (Figure 2 & 3), with binding affinities (Kd) ranging from 9.3-14.3 µM. These Kd values are similar to those calculated for *B. subtilis* Spo0J (56) and below those calculated for *M. xanthus* ParB (55). Based on amino acid sequence alignments and protein structural analyses, there appears to be only a single CTP binding pocket per VirB monomer. Thus, it is likely that the binding stoichiometry of VirB (monomer) to CTP is 1:1. These extrapolations are supported by the ParB literature where the ratio of ParB (monomer) to CTP is also 1:1 (55). Our work demonstrates that VirB is a *bona fide* CTP-binding protein and only the second protein within the fast-evolving clade characterized to bind CTP.

Having established that VirB binds CTP, subsequent alignments with ParB proteins that bind CTP allowed key residues, predicted to lie in the CTP binding pocket of VirB, to be identified. These were located in two separate regions at the N-terminus of VirB (Figure 4A). Residues I65, T68, and F74 (Box I residues (55)) are predicted to directly interact with the cytidine base, whereas G91, R93, and R94 (Box II residues (55)) are predicted to interact with the triphosphate motif of CTP (55–57). In ParB, substitutions of equivalent residues in these two regions led to a direct loss of CTP binding (76, 89) and prevented ParB from undergoing a closed clamp conformation change. These activitiesare necessary for proper partitioning of the DNA (76, 89). In this work, we show that substitution of equivalent residues in VirB leads to a loss of transcriptional anti-silencing activity at the well-characterized VirB-dependent promoter *icsP* (Figure 4B), a loss of the Congo red phenotype, which strongly correlates with *Shigella* virulence (Figure 5) (14), and a loss of discrete focus formation in the *Shigella* cytoplasm when VirB mutants were fused to GFP (Figure 6). Since focus formation of GFP-VirB is wholly dependent on VirB engaging its recognition site *in vivo*, these data provide the first evidence that this cohort of VirB mutants, while stably produced, are defective in forming normal VirB-DNA interactions. Thus, these findings are the first to link the unusual NTP ligand, CTP, to the regulation of *Shigella* virulence genes and hence pathogenicity. Future work will address whether these mutants can (i) bind CTP, (ii) bind DNA, and (iii) how VirB-CTP interactions with the large virulence plasmid impact transcriptional anti-silencing.

Interestingly, no evidence of CTP hydrolysis was observed by VirB in our ITC experiments, although this is an activity demonstrated by the classic ParBs (55–57). A loss of CTP hydrolysis was shown for *M. xanthus* PadC (55), another member of the fast-evolving clade. While two key residues required for hydrolysis, Q127 and K128, are found in VirB (in the so-called Box III, where they are needed for metal coordination and binding the α-phosphate of the NTP, respectively), several members of the fast-evolving clade, including VirB, have a substitution at another highly conserved residue in this region (E to I at position 126 in VirB, which coordinates one of the divalent metal ions needed for CTP hydrolysis). The substitution of this highly conserved acidic residue with a hydrophobic residue could potentially diminish CTPase activity while still supporting CTP-binding (as the other two residues Q127 and K128 are present). This is intriguing for multiple reasons. First, the current ParB model implicates CTP hydrolysis in the reopening of the sliding clamp, thus playing a critical role in the recycling of ParB bound DNA back to the cytoplasmic pool (55–57, 70). Second, the elimination of CTPase activity results in more diffuse ParB-dependent foci in the cytoplasm and defects in chromosome segregation (89, 90), likely due to the inability of ParB to recruit other ParB molecules to distal DNA (91). Hence, the potential lack of CTP hydrolysis in VirB raises the question of how VirB proteins are released from DNA and how VirB forms tight foci without CTPase activity. It is of note that our ITC experiments are performed in the absence of DNA;therefore, CTP hydrolysis may only be triggered when DNA is present. Work is currently underway to address whether VirB is capable of hydrolyzing CTP and whether this activity depends on the presence of DNA.

To conclude, this work provides novel insight into the ParB superfamily through the discovery that there are two distinct clades, the classic ParBs and the fast-evolving clade (Figure 1). We are the first to demonstrate that *Shigella* VirB is a bona fide CTP-binding protein (Figures 2 & 3). With striking differences in both homology and function, VirB is the first plasmid-borne ParB family member shown to bind CTP for its role as a virulence gene regulator. Our findings raise the possibility that other functionally distinct members of the fast-evolving clade may also bind CTP, therefore broadening our understanding of the ParB superfamily, a group of bacterial proteins that play critical roles in many different bacteria. Lastly, our finding that key residues in the predicted CTP binding pocket of VirB, are required for the virulence attributes of *Shigella* (Figures 4, 5, & 6), implicates CTP as an essential ligand for virulence gene regulation in *Shigella* and raises the possibility that virulence may be linked to fluctuations in cellular CTP pools. Future studies that probe how CTP mechanistically influences transcriptional anti-silencing by VirB and that delineate the role CTP plays in the control of *Shigella* virulence are clearly needed.

## Materials and Methods

### Sequence, structure, and phylogenetic analysis

PSIBLAST searches (92) were used to identify homologs of VirB from the NCBI NR database clustered at 90% sequence identity (NR90) by the MMSEQS program (93). To reduce redundancy, the sequences were further clustered with variable length and score thresholds of 0.7 and 0.3 respectively. A multiple sequence alignment was generated using the Mafft program (94) with a local pair algorithm, maxiterate set to 3000, op set to 1.2, and ep set to 0.5.

Secondary structures were predicted using the JPred program (95). Structures were rendered, compared, and superimposed using the Mol* program. Structural models were generated using Alphafold2 program, with templates obtained from HHpred searches for the neural networks deployed by the program.

Phylogenetic analysis was conducted using the maximum likelihood method implemented in the IQtree program (RRID: SCR_017254) (96) with a mixed model combining the Q.pfam substitution matrix, a discrete gamma-distributed model with 4 rate categories and 1 invariant category, and a FreeRate model with 4 rate categories. Bootstrap values were calculated using the ultrafast bootstrap method with 1000 replicates. Phylogenetic trees were rendered using the FigTree program (RRID: SCR_008515) (http://tree.bio.ed.ac.uk/software/figtree/) and rooted via the midpoint rooting. Similar clades were also obtained when the tree was rooted with distant ParB domains such as plant Sulfuredoxins or bacteriophage ParB-like proteins. The differential evolutionary rate plots were calculated using a matrix containing inter-sequence log-corrected pairwise maximum-likelihood distances computed using the FastTree program (97). The distances were centered and scaled using the min-max method and then plotted as Kernel density to normalize the overall area under each curve to 1. A two-sample t-test was used to estimate the significance of the difference between the mean values of the fast-evolving and the slow-evolving clades. All computations were performed using the R language.

### Bacterial strains, plasmids, and media

The strains and plasmids used in this study are listed in Table S1. *S. flexneri* strains were routinely grown on trypticase soy agar (TSA; trypticase soy broth (TSB) containing 1.5% [wt/vol] agar). When appropriate, maintenance of the virulence plasmid was examined by Congo red binding on TSA plates containing 0.01% [wt/vol] Congo red. Depending on the assay, liquid cultures were grown overnight at 30°C in LB broth or minimal media, which limits autofluorescence during imaging (M9 minimal medium supplemented with 0.4% D-glucose, 0.4% Casamino Acids, 0.01 mg/ml nicotinic acid, 0.01 mg/ml tryptophan, 0.01 mg/ml thiamine, 0.1LmM CaCl_2_, and 0.5LmM MgSO_4_) (98, 99). Overnight cultures were diluted 1:100 and subcultured at 37°C with aeration in the specified media (40 mM glycerol replaced D-glucose in minimal medium to allow induction of pBAD to occur (100)). Where necessary, diluted cultures were induced with 0.2% or 0.02% L-arabinose for 3h of a 5h-growth period. To ensure plasmid maintenance, antibiotics were used at the following final concentrations: ampicillin, 100 μg ml^−1^, and chloramphenicol, 25 μg ml^−1^.

### Construction of *virB* mutant alleles

*virB* mutant alleles were created using megaprimer PCR (primers found in Table S2) (101, 102). The resulting alleles were then introduced into pBAD18 using EcoRI and HindIII. Each mutant was diagnosed using EcoRV and verified by Sanger dideoxy sequencing. All constructs made are listed in Table S1 and the primers used are listed in Table S2.

### Construction of plasmids producing GFP-VirB fusions

The *gfp* gene used throughout this work encodes a superfolder GFP (sfGFP) (82, 83). GFP-VirB mutant fusions were produced from pBAD-*sfgfp*-*linker-virB* constructs (81). In the wild-type *virB* coding region of pJNS12 (81), an XhoI site is found at the 5’ end, and a HindIII restriction site is found after the stop codon. Thus, *virB* derivatives (sourced from pATM324 derivatives) were introduced into pJNS12 using these restriction sites to create pBAD-*sfgfp*-*linker-virB* mutant derivatives (pTMG19-21, and pAMO18-19). Each fusion allele was verified by Sanger dideoxy sequencing.

### VirB Protein Purification

VirB was purified by the Monserate Biotechnology Group (San Diego, CA) using a protocol described previously (36).

### Differential Radial Capillary Action of Ligand Assay

Differential radial capillary action of ligand assay, DRaCALA, was adapted from (55–57, 75). Radiolabeled (hot) CTP (5 nM final) and unlabeled (cold) nucleotides (500 µM final) were prepared in buffer containing 500 mM NaCl, 500 mM Tris/HCl pH 7.5, 50 mM MgCl_2_, and kept on ice. Purified VirB protein (50 µM final) was added to each reaction and incubated for 10 minutes at room temperature. After incubation, 4 μL were spotted onto dry nitrocellulose (GE Healthcare) in 15-second increments, allowed to air dry, and then exposed to a storage phosphor screen (Kodak). The screen was imaged on an Amersham Typhoon. Ligand binding was indicated by a darker inner core staining due to rapid immobilization of the protein and was quantified by densitometric analysis of the inner and outer core of each spot using AzureSpot Analysis Software version 2.0.062. The ‘Analysis Toolbox’ was used to draw a circle around the outer and inner core of each spot. Circles were copied using the ‘duplicate’ feature and dragged to adjacent spots so that direct comparisons could be made. The densitometric output for each spot was used as a proxy was used to calculate the fraction bound (F_B_) using the following equation: FB = (I_inner_ - [A_inner_ x [(I_total_ - I_inner_) / (A_total_ - A_inner_)]]) / I_total_ (75).

### Isothermal Titration Calorimetry

The ITC instrument (MicroCal PEAQ-ITC, Malvern Instruments, Worcestershire, United Kingdom) was used at 25°C, with binding buffer consisting of 25 mM HEPES/NaOH, pH 7.6, 100 mM NaCl, 5 mM MgCl_2_, 0.1 mM EDTA, 0.5 mM β-mercaptoethanol, 5% (v/v) glycerol. The measurement cell and injection syringe were washed thoroughly prior to each run. A 300 μL sample of purified VirB protein, at 45-90 μM, or buffer was used to load the measurement cell (200 μL analytical volume). The injection syringe was filled with buffer containing 3 mM of nucleotides as indicated (CTPγS, CTP, or UTP). A single 0.4 µL priming injection was followed by twelve 3.0 µL injections at 150-second intervals. Data were plotted and analyzed using MicroCal PEAQ-ITC Analysis Software to obtain K_d_ values.

### Quantification of transcriptional anti-silencing by VirB derivatives

To quantify the ability of VirB derivatives to regulate transcription, promoter activities of P*icsP-lacZ* (pAFW04) were measured using β-galactosidase assays. Briefly, overnight cultures were diluted in LB containing appropriate antibiotics (ampicillin, 100 μg ml^−1^, or chloramphenicol, 25 μg ml^−1^) and induced, as described previously, these cultures were then lysed and β-galactosidase activity was measured using a modified Miller protocol (11, 99).

### Quantification of protein levels of VirB derivatives

Protein levels of VirB mutants were analyzed by Western Blot analysis, as described in (11, 81). After growth, cells were normalized to cell density (OD_600_), harvested, and washed with 1 mL 0.2 M Tris buffer (pH 8.0). Cells were resuspended in 200 µL 10 mM Tris (pH 7.4), and 50 µL 5X SDS-PAGE buffer. Equal volumes of normalized and heat-denatured protein preparations were electrophoresed on 12.5% SDS-PAGE gels alongside a SeeBlue^TM^ Plus2 Prestained Standard (Invitrogen). VirB was detected using an affinity-purified anti-VirB antibody obtained from Pacific Immunology. A GE anti-rabbit IgG-horseradish peroxidase (HRP; NA9340) secondary antibody was used. All blots were imaged using “auto expose” with chemiluminescence detection (Azure c300; Azure Biosystems).

### Quantification of Congo Red Binding

Congo red binding of each VirB mutant was quantified using the protocol described in reference (100). Briefly, AWY3 (*virB::*Tn5) harboring pBAD, pBAD-*virB,* or pBAD-*virB* mutants were grown overnight in LB supplemented with 0.2% (wt/vol) D-glucose. The following day, cultures were serially diluted (100-fold dilutions) and ∼6µL of each dilution was spotted onto TSA Congo red plates supplemented with either 0.2% (wt/vol) L-arabinose or D-glucose. Plates were incubated overnight at 37°C. Phenotypes were imaged using the auto-exposure setting with a resolution set at 120 µm (Azure 500; Azure Biosystems) using visible light and RGB detection (captured with Cy2 (blue)). The relative amount of Congo red dye bound by the cells was quantified by resuspending three culture spots in 750 µl of 25% ethanol for each sample. Optical densities of cell suspensions were measured at OD_600,_ prior to centrifugation. The OD_498_ of the supernatant was then measured to quantify the amount of Congo red released from cells and normalized to cell density. Relative Congo red bound by cells was calculated using the following equation: [(OD_498_/OD_600_)/ (average (OD_498_/OD_600_)_2457T pBAD_)] × 100.

### Visualization of GFP fusion proteins and quantification of foci

Visualization of GFP fusion proteins and quantification of foci were determined by fluorescence microscopy using the protocol described in (81) with the following exceptions:fluorescence excitation was performed (X-cite 120LED) at a range of 470-525 nm using 50-75% exposure for all captured images and 4-5 fields of view were captured at random for each experimental strain, with roughly 60 cells per field of view. Statistical analyses were performed to compare the distributions of foci as described in statistical analyses.

### Statistical Analyses

Statistical calculations were performed using IBM® SPSS® Statistics for Windows, version 28.01.0. One-way Analysis of Variance (ANOVA) tests were routinely used and post-hoc analyses were performed as indicated in figure legends. Kolmogorov Smirnov test was used to assess statistical differences amongst distributions of foci in VirB mutants. Throughout this work, statistical significance is represented as, *, indicating a *p* < 0.05.

## Supporting information

Supplementary Materials part 1

Supplementary Materials part 2

## Acknowledgments

We thank Monserate Biotechnology Group for the purification of VirB-His_6_, Dr. Tung Le for reagents that were used in proof of principle experiments (data not shown), Dr. Boo Shan Tseng for advice on statistical analyses, and Dr. Marike Palmer for helpful discussions. We thank lab members past and present for insightful discussions. This work was supported by the National Institute of Allergy and Infectious Diseases of the National Institutes of Health (NIH) (R15 AI090573). ITC was supported by the U.S. Army Research Laboratory and the U.S. Army Research Office under grant numbers W911NF-15-1-0043 and W911NF-15-1-0216. UNLV Genomics Core Facility, which was used throughout this study, was supported by the INBRE Program of the National Center for Research Resources [P20 RR-016464]. T.M.G. is a recipient of a U.S. Department of Education GAANN fellowship [P200A210055]. LMI and LA are supported by the intramural funds of the National Library of Medicine, NIH, USA. The content of this paper is solely the responsibility of the authors and does not necessarily represent the official views of NIH. These funding sources had no role in the study design, data collection, and interpretation, or the decision to submit the work for publication.

