## Supplementary Materials part 1 for "VirB, a key transcriptional regulator of *Shigella* virulence, requires a CTP ligand for its regulatory activities"

### Table of Contents

Table S1. Bacterial strains and plasmid used in this study 2

Table S2. Primers used in this study 3

Table S3. Complete statistics for DRaCALA 4

Table S4. Complete statistics for anti-silencing of P*icsP* 5

Table S5. Complete statistics Congo red binding assay 6

Table S6. Complete statistics for fluorescence microscopy distributions 7

Figure S1. Maximum likelihood tree of ParB protein sequences including the VirB-like fast-evolving sequences 8

Figure S2. Multiple sequence alignment of slow-evolving (classic) and fast-evolving (e.g., VirB) ParB proteins containing the catalytic ParB, HTH, tetrahelical and C-terminal domains 9-14

Figure S3. Isothermal Titration Calorimetry buffer-only controls 15

Figure S4. Congo red binding activity of VirB mutants 16

Figure S5. Live cell imaging of GFP-VirB mutants in a *virB* mutant strain of *S. flexneri* 17-19

References 20

**Table S1. Bacterial strains and plasmids used in this study**

| Label | Description | Reference |
|---|---|---|
| **Strains** | | |
| *S. flexneri* | | |
| AWY3 | 2457T *virB*::Tn5; $Kn^r$ | (1) |
| BS103 | 2457T cured of the virulence plasmid | (2) |
| **Plasmids** | | |
| pATM324 | pBAD18-*virB*; $Amp^r$ | (3) |
| pBAD18 | Arabinose-inducible pBAD expression vector, pBR *ori;* $Amp^r$ | (4) |
| pADK15 | pBAD-*virB* K152E; $Amp^r$ | (5) |
| pTMG24 | pBAD-*virB* K152E-R167E; $Amp^r$ | This work |
| pTMG25 | pBAD-*virB* G91S; $Amp^r$ | This work |
| pDRG03 | pBAD-*virB* R93A; $Amp^r$ | This work |
| pDRG04 | pBAD-*virB* R94A; $Amp^r$ | This work |
| pDRG05 | pBAD-*virB* R95A; $Amp^r$ | This work |
| pAMO12 | pBAD-*virB* T68A; $Amp^r$ | This work |
| pAMO13 | pBAD-*virB* T68S; $Amp^r$ | This work |
| pAMO14 | pBAD-*virB* I65A; $Amp^r$ | This work |
| pAMO15 | pBAD-*virB* F74A; $Amp^r$ | This work |
| pAFW04 | pACYC184 carrying WT *virB* in P*icsP-lacZ* | (6) |
| pJNS12 | pBAD-*sfgfp-virB*; $Amp^r$ | (7) |
| pGB682 | pBR322-derived expression vector allowing regulated and dose-dependent recombinant protein expression; *ori* pMB1; $Amp^r$ | (7) |
| pJH66 | pBAD-*linker-sfgfp*; $Amp^r$ | (8) |
| pJNS18 | pBAD-*sfgfp-virB* K152E/R167E; $Amp^r$ | (7) |
| pJNS43 | pBAD-*sfgfp-linker-virB* G91S; $Amp^r$ | This work |
| pTMG19 | pBAD-*sfgfp-linker-virB* R93A; $Amp^r$ | This work |
| pTMG20 | pBAD-*sfgfp-linker-virB* R94A; $Amp^r$ | This work |
| pTMG21 | pBAD-*sfgfp-linker-virB* R95A; $Amp^r$ | This work |
| pAMO18 | pBAD-*sfgfp-linker-virB* T68A; $Amp^r$ | This work |
| pAMO19 | pBAD-*sfgfp-linker-virB* T68S; $Amp^r$ | This work |

**Table S2. Primers used in this study**

| Primer | Sequence 5' to 3' | Description and Use |
|---|---|---|
| W43 | CTCTACTGTTTCTCCATACCC | Sequencing primer for pBAD-*virB* mutants (I65A, T68A, T68S, F74A) |
| W368 | TCATTCGCTAGCAAACCACCCCAATATAAGTTTGAG | Sequencing primer for pBAD-*virB* G91S |
| W453 | AGCGAATTCATAAACAGGGTGTGAT | Sequencing primer used for pBAD-*virB* mutants (R93A, R94A, R95A) |
| W454 | GAAATTCTGGATGGCACTGCTAGAAGAGCATCTGCAATATATGC | Mutagenic primer used to generate pBAD-*virB* R93A |
| W455 | GAAATTCTGGATGGCACTCGTGCAAGAGCATCTGCAATATATGC | Mutagenic primer used to generate pBAD-*virB* R94A |
| W456 | GAAATTCTGGATGGCACTCGTAGAGCAGCATCTGCAATATATGC | Mutagenic primer used to generate pBAD-*virB* R95A |
| W457 | GAGATATTATTTCCTGTGGAACGCTTGC | Primer used to generate megaprimers for pBAD-*virB* mutants (R93A, R94A, R95A); Sequencing primer for pBAD-*virB* mutants (R93A, R94A, R95A) |
| W458 | ATTATTTGCACGGCGTCACACTTTGC | Primer for amplification of pBAD-*virB* derivatives (G91S, R93A. R94A, R95A) |
| W543 | CTAGTCAAAGCTTATGAAGACGATAGATGGCGAGA | Primer for amplification of pBAD-*virB* G91S |
| W563 | CCGAACGAAAAGCGCGACCACATGG | Sequencing primer used for pBAD-*sfgfp-linker-virB* derivatives (T68A, T68S) |
| W638 | TTCTGCGTTCTGATTTAATCTGTATCAGGC | Sequencing primer for pBAD-*virB* mutants (I65A, T68A, T68S, F74A) and pBAD-*sfgfp-linker-virB* derivatives (T68A, T68S); Primer used to amplify pBAD-*virB* DBM |
| W651 | CGAGACAGATTCTCTTTTTTGGCGATATCCTCATAGGACATCCC | Mutagenic primer used to generate pBAD-*virB* K152E |
| W652 | GCACTCGTAGAAGAGCATCTGCA | Primer for amplification of pBAD-*virB* K152E and pBAD-*virB* DBM |
| W653 | GGCTGAAAATCTTCTCTCATCCGCC | Primer for amplification of Box 1 mutants and pBAD-*virB* K152E; Sequencing primer for pBAD-*virB* K152E and DBM |
| W665 | GATTAGCGGATCCTACCTGACGC | Sequencing primer for pBAD-*virB* G91S |
| W721 | TGACTAGCTCGAGGTGGATTTGTGCAACGACTTG | Primer used for amplification of pBAD-*sfgfp-linker-virB* derivatives (T68A, T68S) |
| W722 | GTATATCGTTTGCTAGTTTTCTGGC | Primer used for amplification of pBAD-*sfgfp-linker-virB* T68A and T68S |
| W752 | TGCTGCCTGAAAGGCCTCAGTGACTTTCGCGCGAGACAG | Mutagenic primer used to generate pBAD-*virB* DBM |
| W841 | TACGAACCTGTAATAGGAAGGGAGATTGATGGTAGAATTGAAATTCTGGATAGCACTCGT | Mutagenic primer for amplification of pBAD-*virB* G91S |
| W900 | CATCATCATCATGGTATGGCTAGCG | Primer used to generate megaprimer for pBAD-*virB* (I65A,T68A, T68S, F74A) |
| W901 | GAATTGTTGTAGCTTTATAGCTTTTATGATATCGGC | Primer for amplification of pBAD-*virB* T68A |
| W902 | CAGAAAATTAAGACCAATACCAAGTTCTCGG | Sequencing primer used for pBAD-*virB* mutants (I65A, T68A, T68S, F74A) |
| W921 | GAATTGTTGTAGCTTTATAGATTTTATGATATCGGC | Primer for amplification of pBAD-*virB* T68S |
| W925 | GCTTTATAGTTTTTATGGCATCGGCTAGTG | Primer for amplification of pBAD-*virB* I65A |
| W926 | CTATTACAGGGAAGGCTTGTTGTAGC | Primer for amplification of pBAD-*virB* F74A |

**Table S3. Complete statistics for DRaCALA competition assay**

| Competition Assay | | no protein | VirB + cold competitor | | | | |
|---|---|---|---|---|---|---|---|
| | | | no cold | CTP | UTP | ATP | GTP |
| no protein | | | <0.001 * | <0.001 * | <0.001 * | <0.001 * | <0.001 * |
| VirB + cold competitor | no cold | | | <0.001 * | 0.054 | 0.983 | 0.063 |
| | CTP | | | | <0.001 * | <0.001 * | <0.001 * |
| | UTP | | | | | 1 | 1 |
| | ATP | | | | | | 1 |
| | GTP | | | | | | |

Significance was determined using a one-way ANOVA with post hoc Bonferroni. Asterisks indicate $p<0.05$. Grey boxes represent data that was not compared.

**Table S4. Complete statistics for anti-silencing of P*icsP***

| VirB Anti-silencing Activity | WT | Empty | K152E | DBM | G91S | R93A | R94A | R95A | T68A | T68S | I65A | F74A |
|---|---|---|---|---|---|---|---|---|---|---|---|---|
| **WT** | | <0.001 * | <0.001 * | <0.001 * | <0.001 * | <0.001 * | <0.001 * | <0.001 * | <0.001 * | 1 | <0.001 * | <0.001 * |
| **Empty** | | | 0.593 | 1 | 1 | 1 | 1 | <0.001 * | 0.998 | <0.001 * | 1 | 1 |
| **K152E** | | | | 0.825 | 0.798 | 0.470 | 0.693 | <0.001 * | 0.989 | <0.001 * | 0.766 | 0.778 |
| **DBM** | | | | | 1 | 1 | 1 | <0.001 * | 1 | <0.001 * | 1 | 1 |
| **G91S** | | | | | | 1 | 1 | <0.001 * | 1 | <0.001 * | 1 | 1 |
| **R93A** | | | | | | | 1 | <0.001 * | 0.989 | <0.001 * | 1 | 1 |
| **R94A** | | | | | | | | <0.001 * | 1 | <0.001 * | 1 | 1 |
| **R95A** | | | | | | | | | <0.001 * | <0.001 * | <0.001 * | <0.001 * |
| **T68A** | | | | | | | | | | <0.001 * | 1 | 1 |
| **T68S** | | | | | | | | | | | <0.001 * | <0.001 * |
| **I65A** | | | | | | | | | | | | 1 |
| **F74A** | | | | | | | | | | | | |

Significance was determined using a two-way ANOVA with post hoc Tukey HSD. Asterisks indicate p<0.05. Grey boxes represent data that was not compared.

**Table S5. Complete statistics for Congo red binding assay**

| Congo Red Binding Activity (0.2% L-ara) | WT | Empty | K152E | DBM | R93A | R94A | R95A | T68A | T68S | I65A | F74A |
|---|---|---|---|---|---|---|---|---|---|---|---|
| WT | | <0.001 * | <0.001 * | <0.001 * | <0.001 * | <0.001 * | 0.002* | <0.001 * | 0.759 | <0.001 * | <0.001 * |
| Empty | | | <0.001 * | 1 | 1 | 1 | <0.001 * | 0.993 | <0.001 * | 1 | 1 |
| K152E | | | | <0.001 * | <0.001 * | <0.001 * | <0.001 * | <0.001 * | 0.010* | <0.001 * | <0.001 * |
| DBM | | | | | 1 | 1 | <0.001 * | 0.999 | <0.001 * | 1 | 1 |
| R93A | | | | | | 1 | <0.001 * | 0.995 | <0.001 * | 1 | 1 |
| R94A | | | | | | | <0.001 * | 0.999 | <0.001 * | 1 | 1 |
| R95A | | | | | | | | <0.001 * | <0.001 * | <0.001 * | <0.001 * |
| T68A | | | | | | | | | <0.001 * | 1 | 0.999 |
| T68S | | | | | | | | | | <0.001 * | <0.001 * |
| I65A | | | | | | | | | | | 1 |
| F74A | | | | | | | | | | | |

| Congo Red Binding Activity (0.2% Glu) | WT | Empty | K152E | DBM | R93A | R94A | R95A | T68A | T68S | I65A | F74A |
|---|---|---|---|---|---|---|---|---|---|---|---|
| WT | | 0.853 | 1 | 1 | 1 | 1 | 1 | 1 | 1 | 0.999 | 1 |
| Empty | | | 0.991 | 0.988 | 0.984 | 0.967 | 0.957 | 0.946 | 0.947 | 0.999 | 0.976 |
| K152E | | | | 1 | 1 | 1 | 1 | 1 | 1 | 1 | 1 |
| DBM | | | | | 1 | 1 | 1 | 1 | 1 | 1 | 1 |
| R93A | | | | | | 1 | 1 | 1 | 1 | 1 | 1 |
| R94A | | | | | | | 1 | 1 | 1 | 1 | 1 |
| R95A | | | | | | | | 1 | 1 | 1 | 1 |
| T68A | | | | | | | | | 1 | 1 | 1 |
| T68S | | | | | | | | | | 1 | 1 |
| I65A | | | | | | | | | | | 1 |
| F74A | | | | | | | | | | | |

Significance was determined using a two-way ANOVA with post hoc Tukey HSD. Asterisks indicate p<0.05. Grey boxes represent data that was not compared.

**Table S6. Complete statistics for fluorescence microscopy distributions**

| Focus Formation | WT | Empty | GFP | DBM | R93A | R94A | R95A | T68A | T68S |
|---|---|---|---|---|---|---|---|---|---|
| WT | | <.001 * | <.001 * | <.001 * | <.001 * | <.001 * | <.001 * | <.001 * | .054 |
| Empty | | | <.001 * | <.001 * | <.001 * | <.001 * | <.001 * | <.001 * | <.001 * |
| GFP | | | | <.001 * | <.001 * | <.001 * | <.001 * | .147 | <.001 * |
| DBM | | | | | .965 | .190 | 0.099 | <.001 * | <.001 * |
| R93A | | | | | | .211 | 0.537 | <.001 * | <.001 * |
| R94A | | | | | | | 0.118 | .024* | <.001 * |
| R95A | | | | | | | | <.001 * | <.001 * |
| T68A | | | | | | | | | <.001 * |
| T68S | | | | | | | | | |

Significance was determined using a Kolmogorov-Smirnov test with post hoc Bonferroni. Asterisks indicate $p<0.05$. Grey boxes represent data that was not compared.

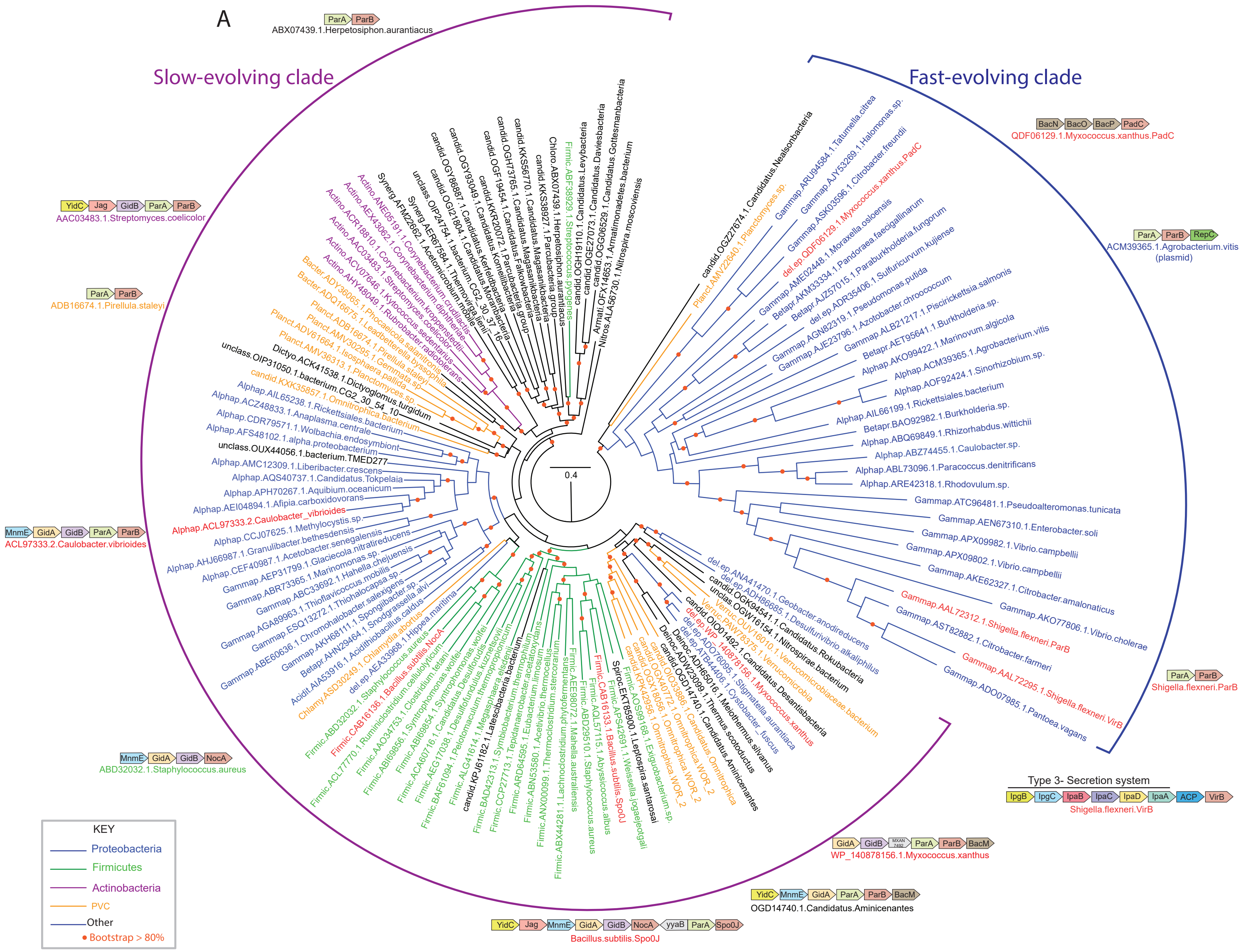


**Figure S1. Full maximum likelihood tree of ParB protein sequences including the VirB-like fast-evolving sequences.** (A) Maximum likelihood tree of ParB protein sequences including the VirB-like fast-evolving sequences. The tree was computed using IQtree and bootstrapped with 1000 replicates. Nodes and branches corresponding to widely represented phylogenetic clades are colored according to the key. Protein sequences are denoted with their clade, gene and species name. The slow- and fast-evolving clades are marked and representative gene neighborhoods of distinct clades of the tree are shown. Genes in gene neighborhoods are shown as box arrows with the arrowhead pointing to the gene at the 3' end. Clade abbreviations include: Acidit: Acidiothiobacillus, Actino: Actinomycetes, Alphap: Alpha-proteobacteria, Armati: Armatimonas, Bacter: Bacteroidetes, Betapr: Beta-proteobacteria, candid: Dark-matter bacteria, Chlamy: Chlamydiae, Chloro: Chloroflexi, Deinoc: Thermus/Deinococcus, del.ep: Delta/epsilon proteobacteria, Dictyo: Dictyoglomus, Firmic: Firmicutes, Gammap: Gamma-proteobacteria, Nitros: Nitrospirae, Planct: Planctomycetes, Spiroc: Spirochaetes, Synerg: Synergistetes, Verruc: Verrucomicrobiae, unclass: Unclassified bacteria.
