## Supplementary Materials part 2 for "VirB, a key transcriptional regulator of *Shigella* virulence, requires a CTP ligand for its regulatory activities"

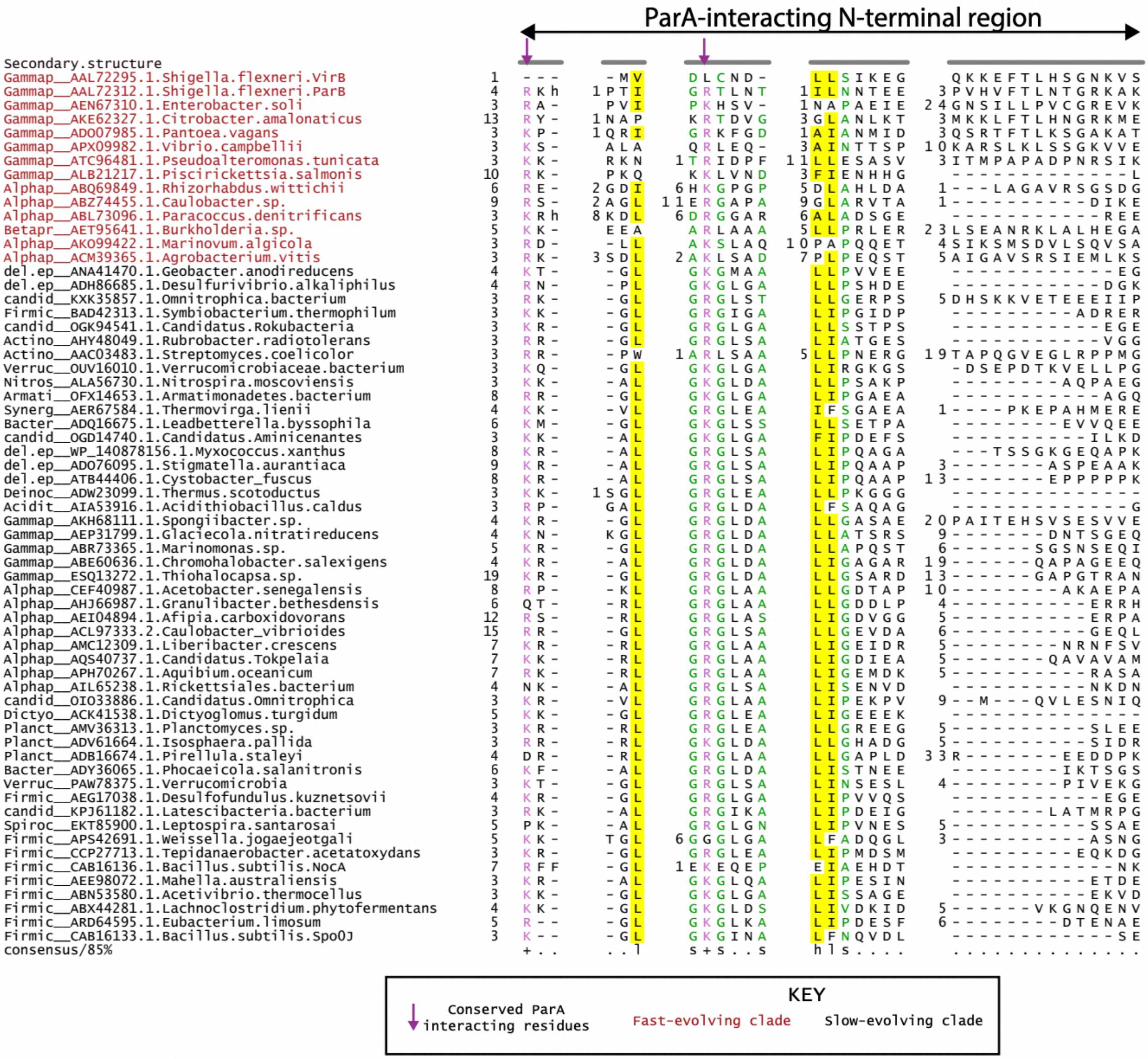
ParA-interacting N-terminal region
Secondary.structure
consensus/85%
KEY
Conserved ParA interacting residues
Fast-evolving clade
Slow-evolving clade

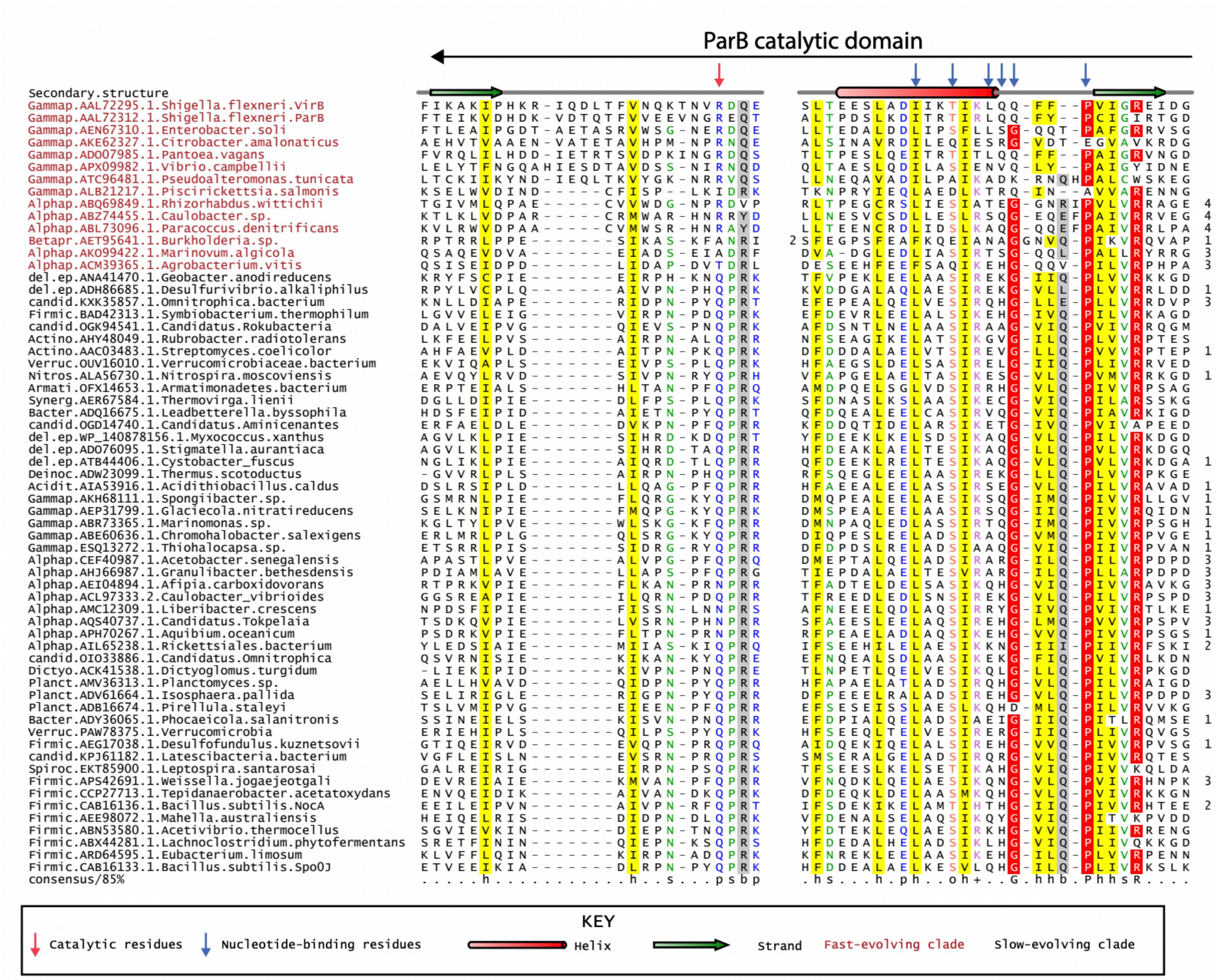
ParB catalytic domain
Secondary.structure
Gammap.AAL72295.1.Shigella.flexneri.VirB
Gammap.AAL72312.1.Shigella.flexneri.ParB
Gammap.AEN67310.1.Enterobacter.soli
Gammap.AKE62327.1.Citrobacter.amalonaticus
Gammap.ADO07985.1.Pantoea.vagans
Gammap.APX09982.1.Vibrio.campbellii
Gammap.ATC96481.1.Pseudoalteromonas.tunicata
Gammap.ALB21217.1.Piscirickettsia.salmonis
Alphap.ABQ69849.1.Rhizorhabdus.wittichii
Alphap.ABZ74455.1.Caulobacter.sp.
Alphap.ABL73096.1.Paracoccus.denitrificans
Betapr.AET95641.1.Burkholderia.sp.
Alphap.AKO99422.1.Marinovum.algicola
Alphap.ACM39365.1.Agrobacterium.vitis
consensus/85%
KEY
Catalytic residues
Nucleotide-binding residues
Helix
Strand
Fast-evolving clade
Slow-evolving clade

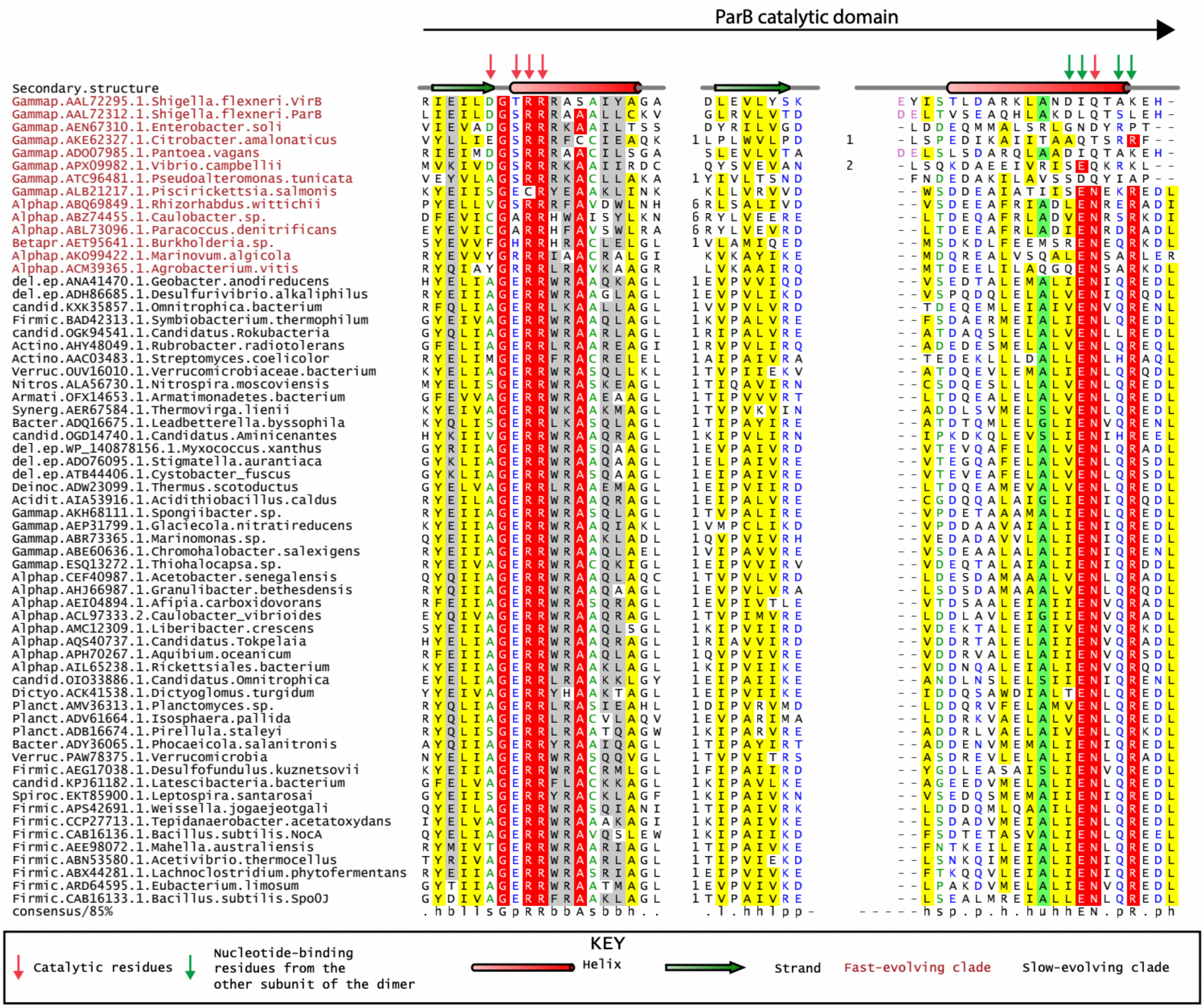

ParB catalytic domain
Secondary.structure
Gammap.AAL72295.1.Shigella.flexneri.VirB
Gammap.AAL72312.1.Shigella.flexneri.ParB
Gammap.AEN67310.1.Enterobacter.soli
Gammap.AKE62327.1.Citrobacter.amalonaticus
Gammap.ADO07985.1.Pantoea.vagans
Gammap.APX09982.1.Vibrio.campbellii
Gammap.ATC96481.1.Pseudoalteromonas.tunicata
Gammap.ALB21217.1.Piscirickettsia.salmonis
Alphap.ABQ69849.1.Rhizorhabdus.wittichii
Alphap.ABZ74455.1.Caulobacter.sp.
Alphap.ABL73096.1.Paracoccus.denitrificans
Betapr.AET95641.1.Burkholderia.sp.
Alphap.AKO99422.1.Marinovum.algicola
Alphap.ACM39365.1.Agrobacterium.vitis
del.ep.ANA41470.1.Geobacter.anodireducens
del.ep.ADH86685.1.Desulfurivibrio.alkaliphilus
candid.KXK35857.1.Omnitrophica.bacterium
Firmic.BAD42313.1.Symbiobacterium.thermophilum
candid.OGK94541.1.Candidatus.Rokubacteria
Actino.AHY48049.1.Rubrobacter.radiotolerans
Actino.AAC03483.1.Streptomyces.coelicolor
Verruc.OUV16010.1.Verrucomicrobiaceae.bacterium
Nitros.ALA56730.1.Nitrospira.moscoviensis
Armati.OFX14653.1.Armatimonadetes.bacterium
Synerg.AER67584.1.Thermovirga.lienii
Bacter.ADQ16675.1.Leadbetterella.byssophila
candid.OGD14740.1.Candidatus.Aminicenantes
del.ep.WP_140878156.1.Myxococcus.xanthus
del.ep.ADO76095.1.Stigmatella.aurantiaca
del.ep.ATB44406.1.Cystobacter_fuscus
Deinoc.ADW23099.1.Thermus.scotoductus
Acidit.AIA53916.1.Acidithiobacillus.caldus
Gammap.AKH68111.1.Spongiibacter.sp.
Gammap.AEP31799.1.Glaciecola.nitratireducens
Gammap.ABR73365.1.Marinomonas.sp.
Gammap.ABE60636.1.Chromohalobacter.salexigens
Gammap.ESQ13272.1.Thiohalocapsa.sp.
Alphap.CEF40987.1.Acetobacter.senegalensis
Alphap.AHJ66987.1.Granulibacter.bethesdensis
Alphap.AEI04894.1.Afipia.carboxidovorans
Alphap.ACL97333.2.Caulobacter_vibrioides
Alphap.AMC12309.1.Liberibacter.crescens
Alphap.AQS40737.1.Candidatus.Tokpelaia
Alphap.APH70267.1.Aquibium.oceanicum
Alphap.AIL65238.1.Rickettsiales.bacterium
candid.OIO33886.1.Candidatus.Omnitrophica
Dictyo.ACK41538.1.Dictyoglomus.turgidum
Planct.AMV36313.1.Planctomyces.sp.
Planct.ADV61664.1.Isosphaera.pallida
Planct.ADB16674.1.Pirellula.staleyi
Bacter.ADY36065.1.Phocaeicola.salanitronis
Verruc.PAW78375.1.Verrucomicrobia
Firmic.AEG17038.1.Desulfofundulus.kuznetsovii
candid.KPJ61182.1.Latescibacteria.bacterium
Spiroc.EKT85900.1.Leptospira.santarosai
Firmic.APS42691.1.Weissella.jogaejeotgali
Firmic.CCP27713.1.Tepidanaerobacter.acetatoxydans
Firmic.CAB16136.1.Bacillus.subtilis.NocA
Firmic.AEE98067.1.Mahella.australiensis
Firmic.ABN53580.1.Acetivibrio.thermocellus
Firmic.ABX44281.1.Lachnoclostridium.phytofermentans
Firmic.ARD64595.1.Eubacterium.limosum
Firmic.CAB16133.1.Bacillus.subtilis.Spo0J
consensus/85%
KEY
Catalytic residues
Nucleotide-binding residues from the other subunit of the dimer
Helix
Strand
Fast-evolving clade
Slow-evolving clade

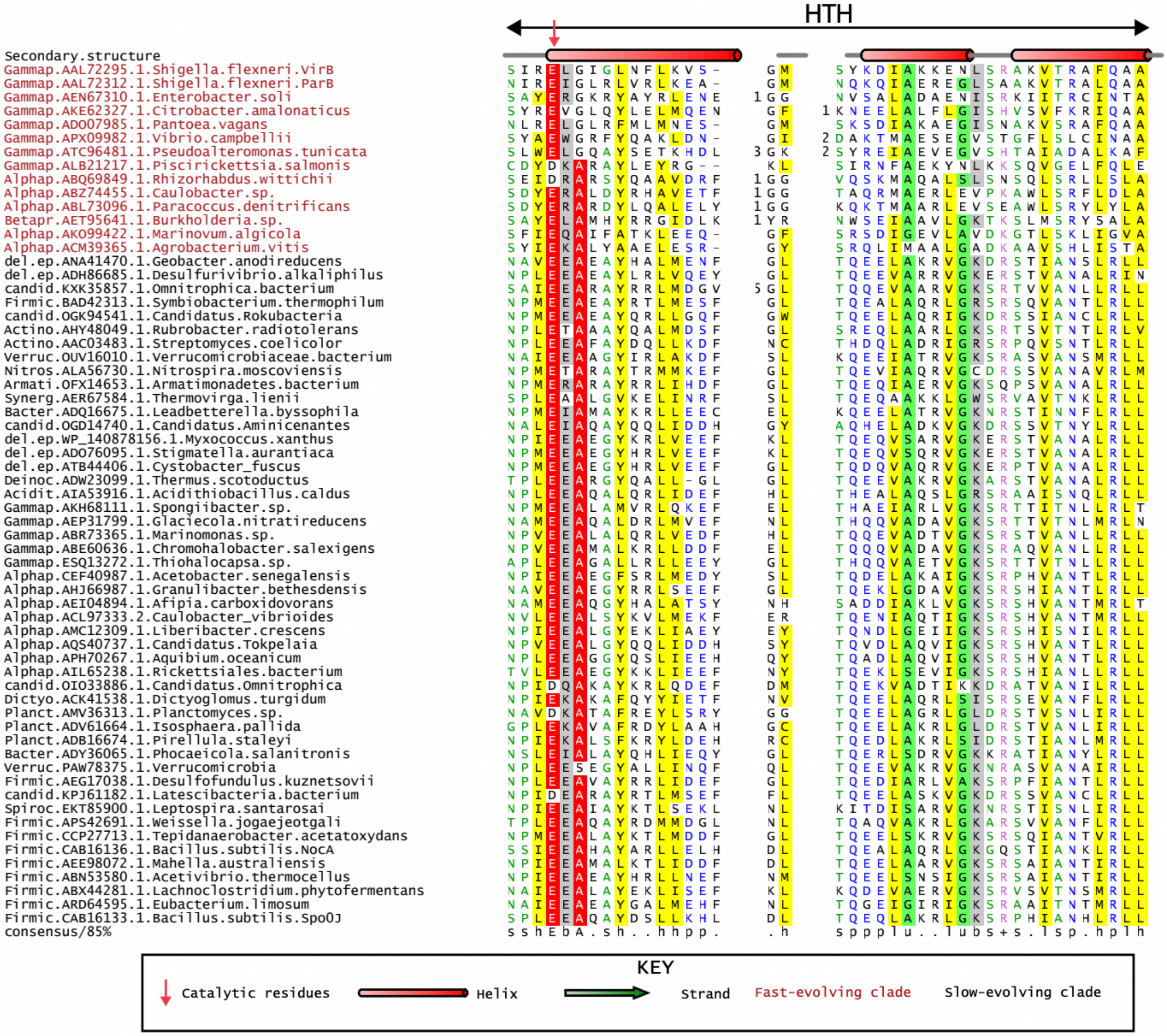
HTH
Secondary.structure
Gammap.AAL72295.1.Shigella.flexneri.VirB
Gammap.AAL72312.1.Shigella.flexneri.ParB
Gammap.AEN67310.1.Enterobacter.soli
Gammap.AKE62327.1.Citrobacter.amalonaticus
Gammap.ADO07985.1.Pantoea.vagans
Gammap.APX09982.1.Vibrio.campbellii
Gammap.ATC96481.1.Pseudoalteromonas.tunicata
Gammap.ALB21217.1.Piscirickettsia.salmonis
Alphap.ABQ69849.1.Rhizorhabdus.wittichii
Alphap.ABZ74455.1.Caulobacter.sp.
Alphap.ABL73096.1.Paracoccus.denitrificans
Betapr.AET95641.1.Burkholderia.sp.
Alphap.AKO99422.1.Marinovum.algicola
Alphap.ACM39365.1.Agrobacterium.vitis
del.ep.ANA41470.1.Geobacter.anodireducens
del.ep.ADH86685.1.Desulfurivibrio.alkaliphilus
candid.KXK35857.1.Omnitrophica.bacterium
Firmic.BAD42313.1.Symbiobacterium.thermophilum
candid.OGK94541.1.Candidatus.Rokubacteria
Actino.AHY48049.1.Rubrobacter.radiotolerans
Actino.AAC03483.1.Streptomyces.coelicolor
Verruc.OUV16010.1.Verrucomicrobiaceae.bacterium
Nitros.ALA56730.1.Nitrospira.moscoviensis
Armati.OFX14653.1.Armatimonadetes.bacterium
Synerg.AER67584.1.Thermovirga.lienii
Bacter.ADQ16675.1.Leadbetterella.byssophila
candid.OGD14740.1.Candidatus.Aminicenantes
del.ep.WP_140878156.1.Myxococcus.xanthus
del.ep.ADO76095.1.Stigmatella.aurantiaca
del.ep.ATB44406.1.Cystobacter_fuscus
Deinoc.ADW23099.1.Thermus.scotoductus
Acidit.AIA53916.1.Acidithiobacillus.caldus
Gammap.AKH68111.1.Spongiibacter.sp.
Gammap.AEP31799.1.Glaciecola.nitratireducens
Gammap.ABR73365.1.Marinomonas.sp.
Gammap.ABE60636.1.Chromohalobacter.salexigens
Gammap.ESQ13272.1.Thiohalocapsa.sp.
Alphap.CEF40987.1.Acetobacter.senegalensis
Alphap.AHJ66987.1.Granulibacter.bethesdensis
Alphap.AEI04894.1.Afipia.carboxidovorans
Alphap.ACL97333.2.Caulobacter_vibrioides
Alphap.AMC12309.1.Liberibacter.crescens
Alphap.AQS40737.1.Candidatus.Tokpelaia
Alphap.APH70267.1.Aquibium.oceanicum
Alphap.AIL65238.1.Rickettsiales.bacterium
candid.OIO33886.1.Candidatus.Omnitrophica
Dictyo.ACK41538.1.Dictyoglomus.turgidum
Planct.AMV36313.1.Planctomyces.sp.
Planct.ADV61664.1.Isosphaera.pallida
Planct.ADB16674.1.Pirellula.staleyi
Bacter.ADY36065.1.Phocaeicola.salanitronis
Verruc.PAW78375.1.Verrucomicrobia
Firmic.AEG17038.1.Desulfofundulus.kuznetsovii
candid.KPJ61182.1.Latescibacteria.bacterium
Spiroc.EKT85900.1.Leptospira.santarosai
Firmic.APS42691.1.Weissella.jogaejeotgali
Firmic.CCP27713.1.Tepidanaerobacter.acetatoxydans
Firmic.CAB16136.1.Bacillus.subtilis.NocA
Firmic.AEE98072.1.Mahella.australiensis
Firmic.ABN53580.1.Acetivibrio.thermocellus
Firmic.ABX44281.1.Lachnoclostridium.phytofermentans
Firmic.ARD64595.1.Eubacterium.limosum
Firmic.CAB16133.1.Bacillus.subtilis.Spo0J
consensus/85%
KEY
Catalytic residues
Helix
Strand
Fast-evolving clade
Slow-evolving clade

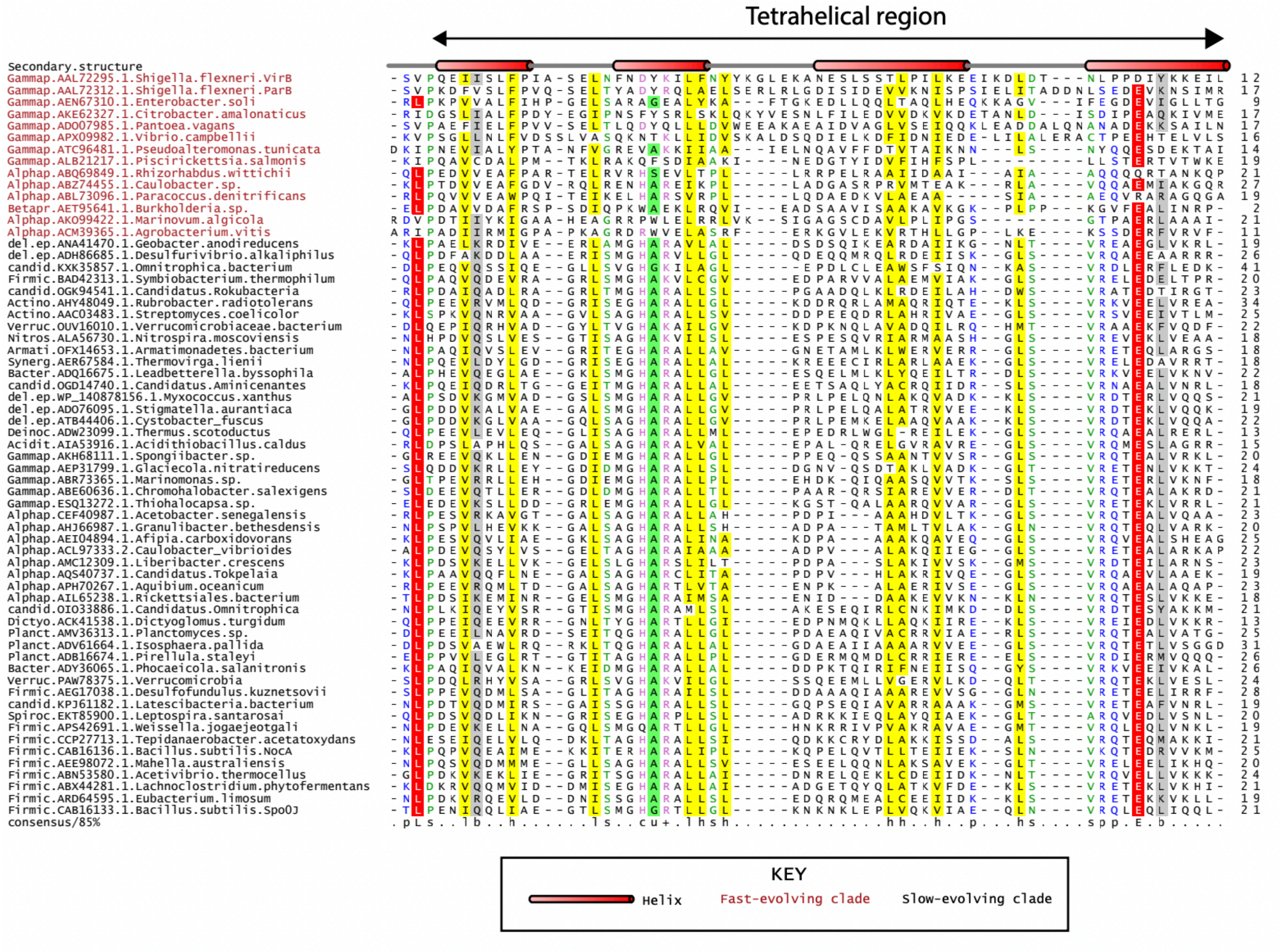
Tetrahelical region
Secondary.structure
consensus/85%
KEY
Helix
Fast-evolving clade
Slow-evolving clade

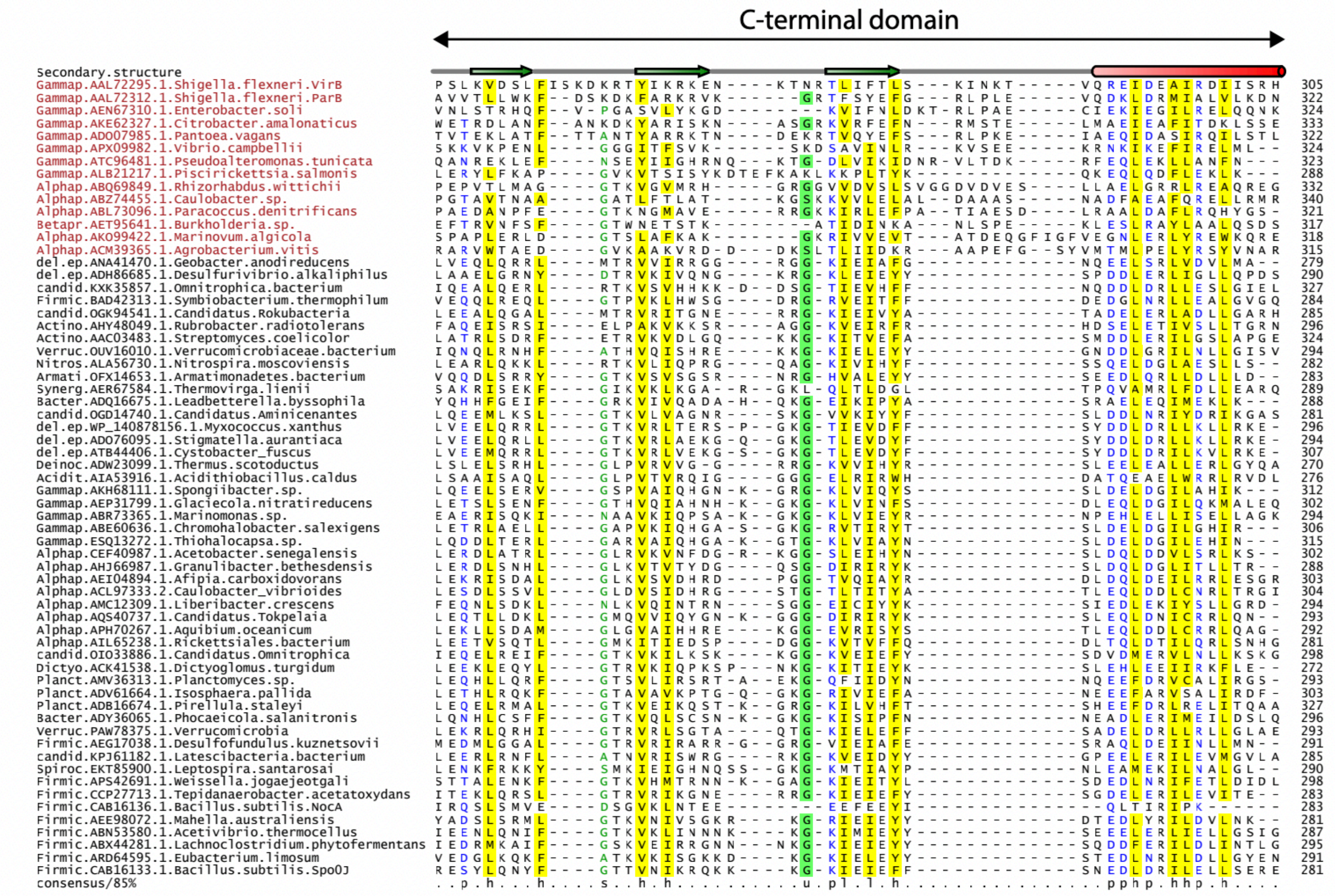


**Figure S2. Multiple sequence alignment of slow-evolving (classic) and fast-evolving (e.g., VirB) ParB proteins containing the catalytic ParB, HTH, tetrahelical and C-terminal domains.** Taxonomic clade name, Genbank accession, and species name separated by dots are denoted. Secondary structure and the sequence consensus at 85% identity are depicted above and below the alignment, respectively. Coloring of sequence columns is as per the residue consensus abbreviation. Consensus abbreviations and coloring scheme are as follows: h: hydrophobic (FWYILVACM), a: aromatic (FWY) and l: aliphatic (ILV) residues shaded yellow; c: charged (DEHKR) and +: basic (KRH) residues colored magenta, b: big (LIFMWYERKQ) residues shaded grey, o: alcohol-group (ST) residues colored red, p: polar (STECDRKHNQ) residues colored blue, s: small (AGSCDNPTV) residues colored green, and u: tiny (GAS) residues shaded green. Fully conserved residues are shaded red. α-helices and β-strands are depicted as cylinders and arrows, respectively. Numbers between aligned blocks indicate the number of poorly conserved residues that were omitted from the alignment for brevity. Predicted catalytic and nucleotide-interacting positions are indicated. Taxonomic clade abbreviations are as follows; Acidit: Acidithiobacillus, Actino: Actinobacteria, Armati: Armatimonadetes, Bacter: Bacteroidetes, candid: Candidatus (uncultured bacteria), Deinoc: Thermus/Deinococcus, Dictyo: Dictyoglomus, Firmic: Firmicutes, Nitros: Nitrospira, Planct: Planctomycetes, Alphap: Alphaproteobacteria, Betapr: Betaproteobacteria, Gammp: Gamma proteobacteria, del.ep: delta/epsilon protebacteria, Spiroc: Spirochaetes, Synerg: Synergistes, Verruc: Verrucomicrobiae

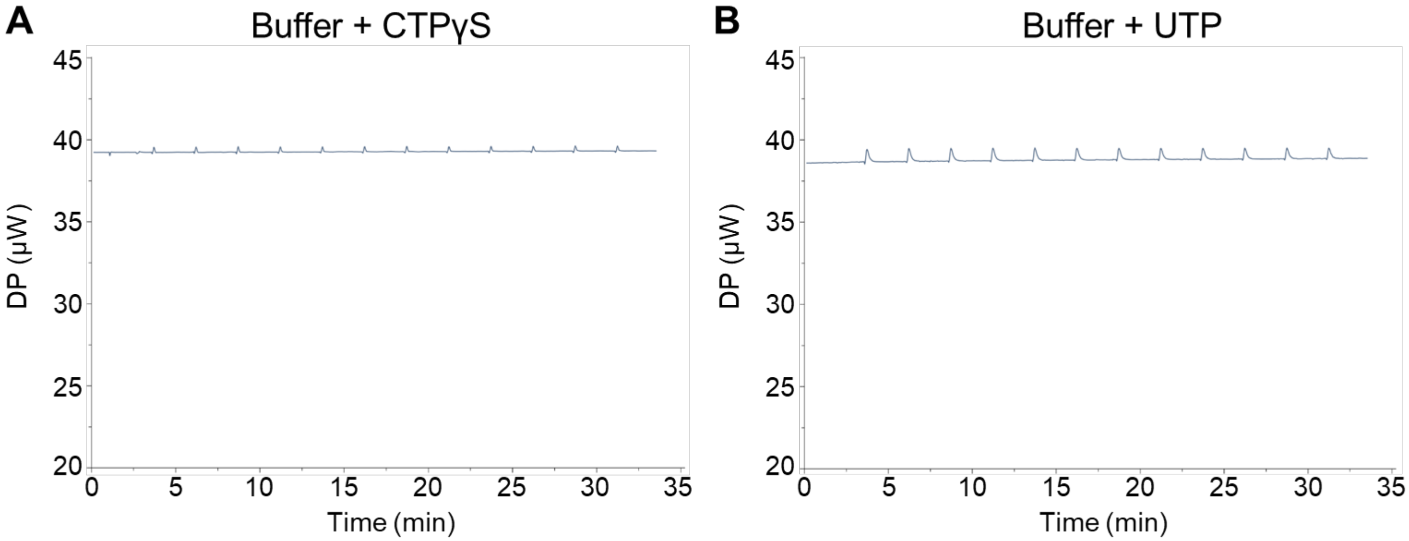


**Figure S3. Isothermal Titration Calorimetry buffer-only controls.** ITC Measurements of (A) VirB buffer-only and 3 mM CTPγS in the presence of $Mg^{2+}$, and (B) VirB buffer-only and 3 mM UTP in the presence of $Mg^{2+}$.

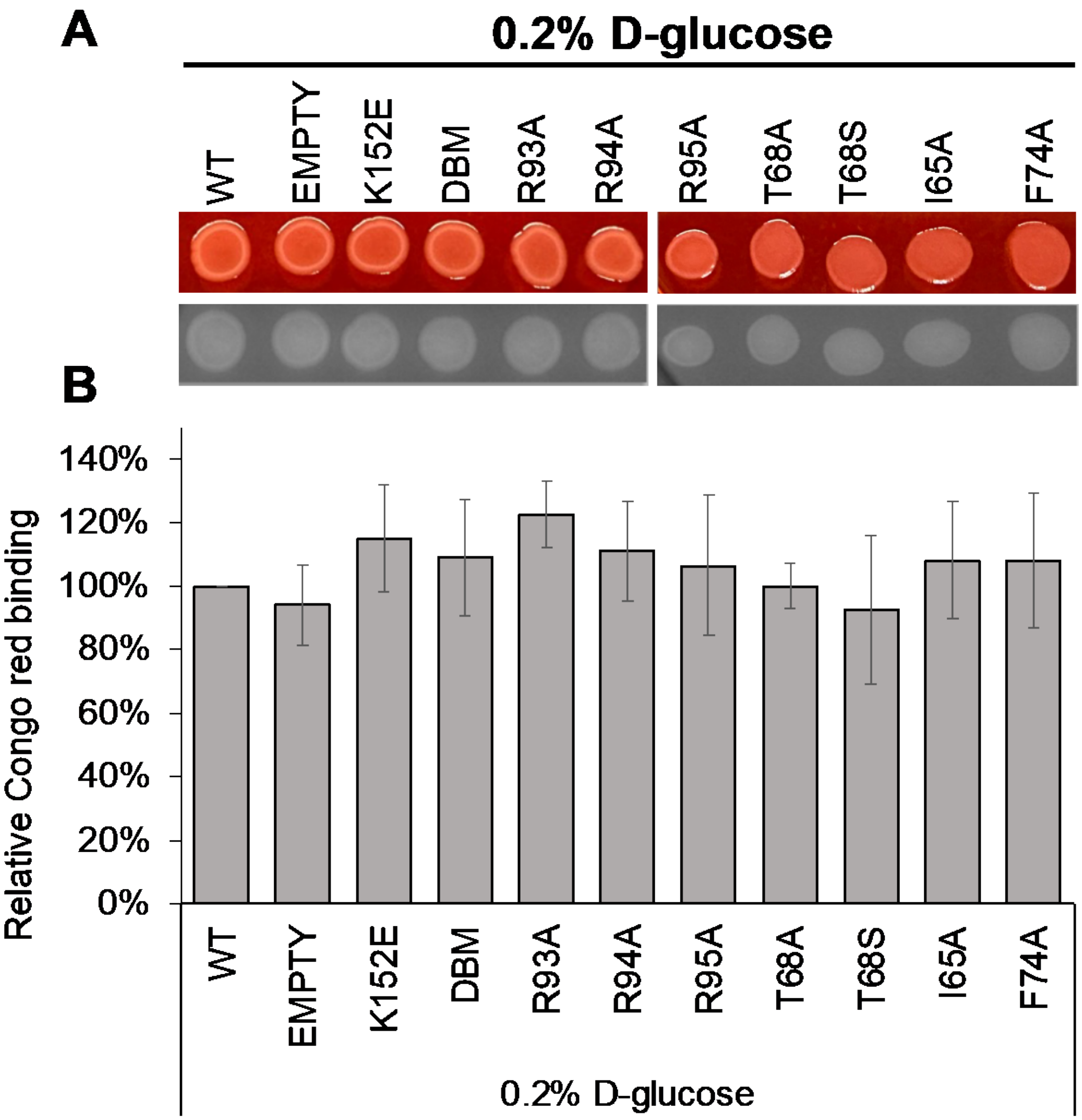


**Figure S4. Congo red binding activity of VirB mutants.** (A) Congo red binding by *S. flexneri virB*::Tn5 harboring pBAD-VirB derivatives under non-inducing conditions. Images were captured using visible light (top) and blue light (Cy2) (bottom). (B) Quantitative analysis of Congo red binding *S. flexneri virB*::Tn5 harboring pBAD-VirB derivatives (non-induced). Relative Congo red binding was calculated as $[(OD_{498}/OD_{600})/\ (\text{average}\ (OD_{498}/OD_{600})_{2457T\ pBAD})] \times 100$. Assays were completed with three biological replicates and repeated three times. Representative data are shown. Significance was calculated using a one-way ANOVA with post hoc Tukey HSD, $p < 0.05$. *, statistically significant compared to wild-type. Complete statistical analysis is provided in Supplementary Table S5.

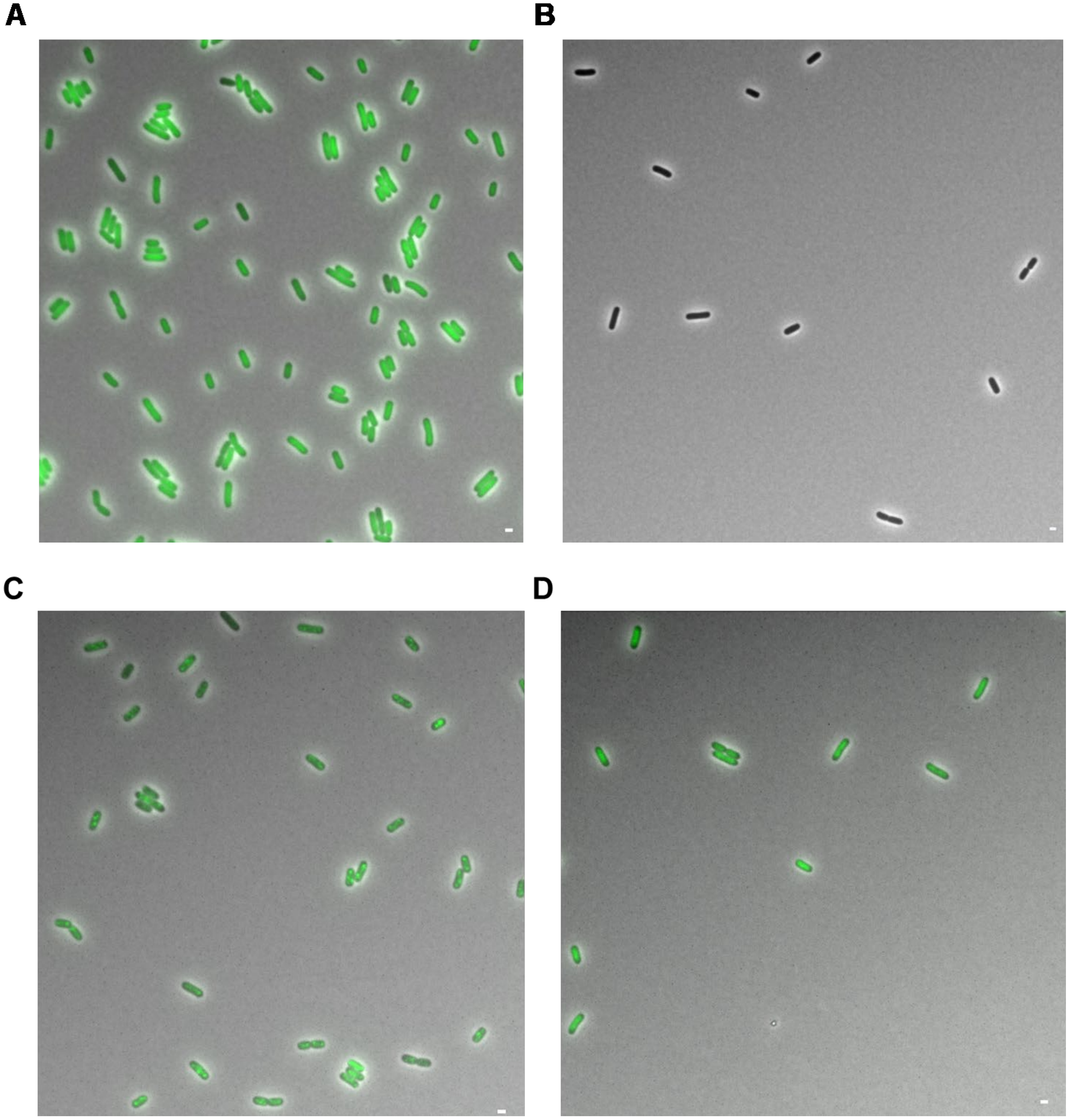
A
B
C
D

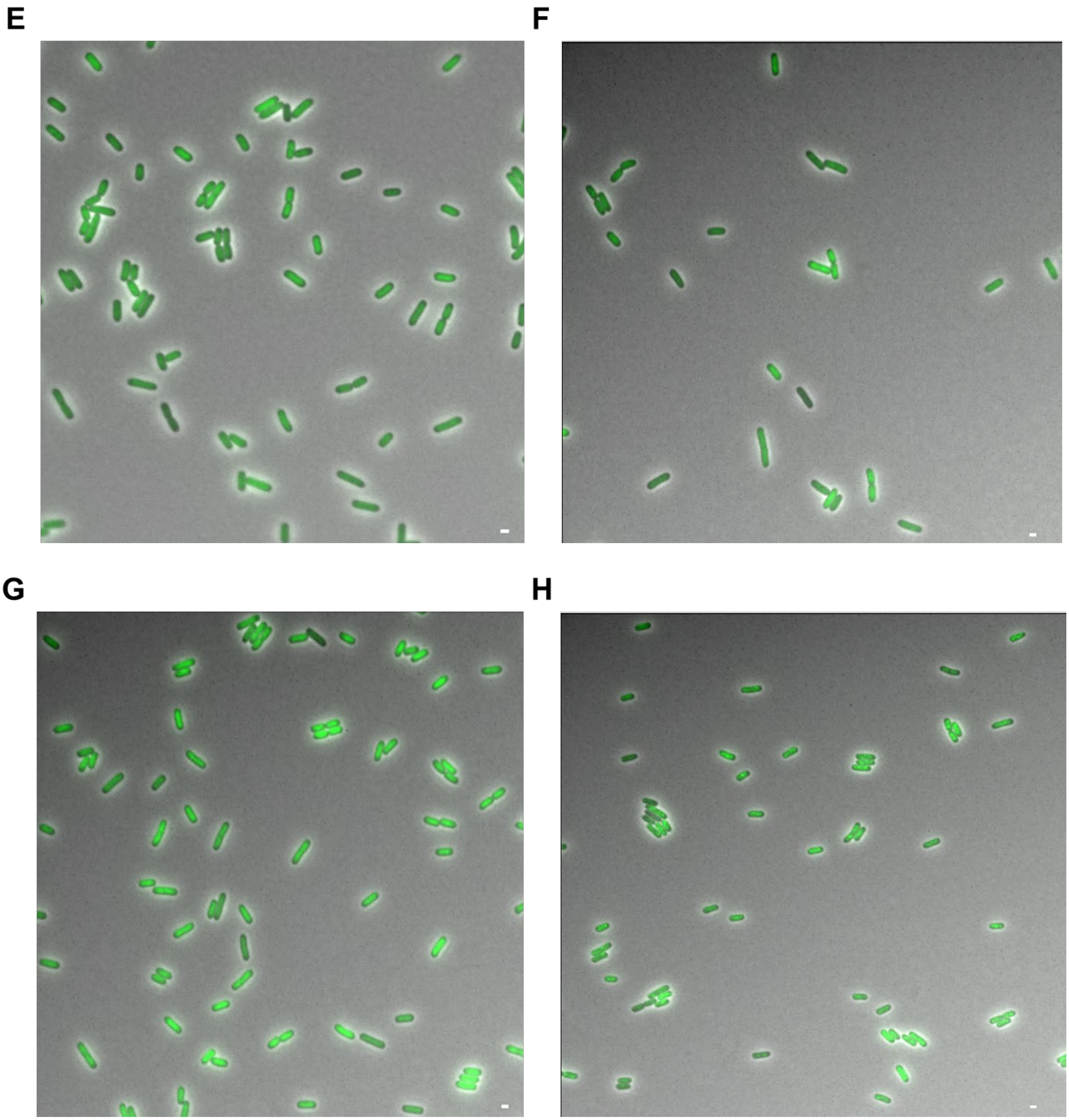
E
F
G
H

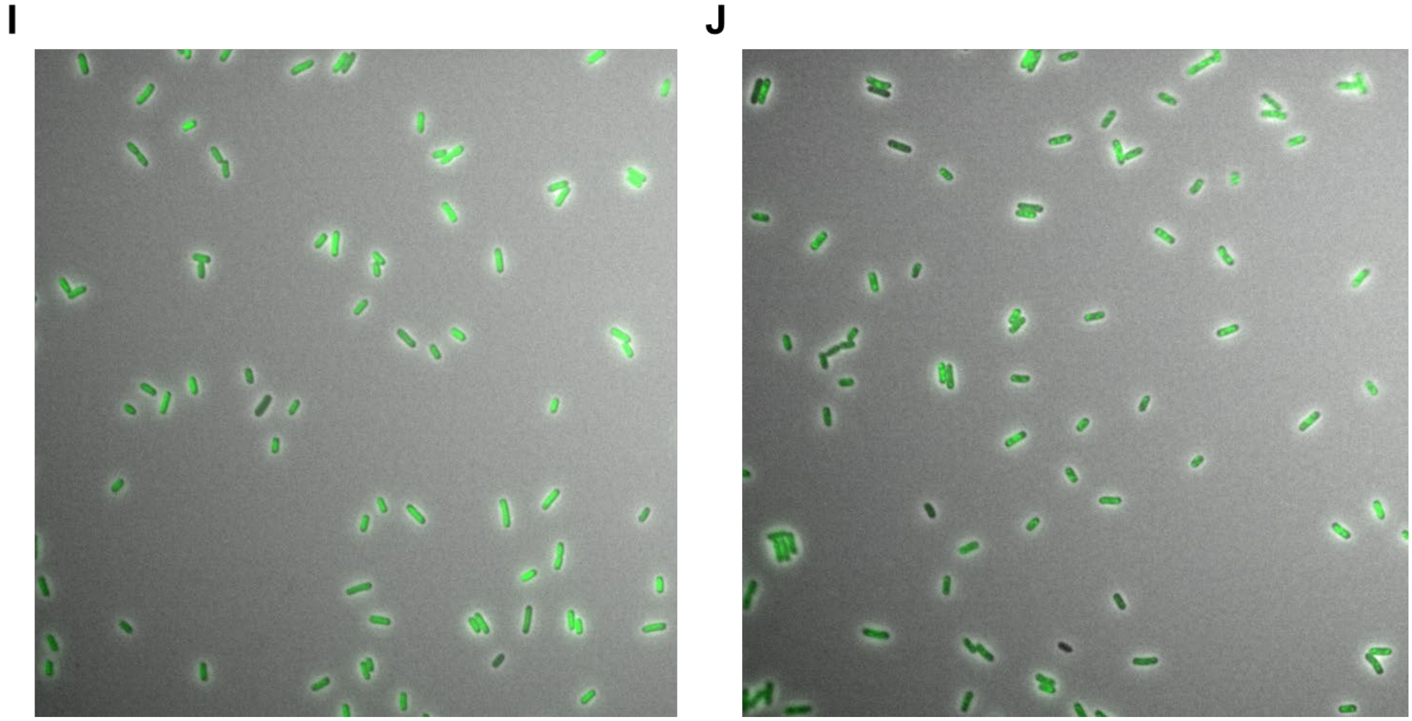


**Figure S5. Live cell imaging of GFP-VirB mutants in a *virB* mutant strain of *S. flexneri*.** Representative fields of view show either nucleoid-associated or diffuse signals for (A) GFP, (B) EMPTY, (C) GFP-VirB, (D) GFP-VirB DBM, (E) GFP-VirB G91S, (F) GFP-VirB R93A, (G) GFP-VirB R94A, (H) GFP-VirB R95A, (I) GFP-VirB T68A, (J) GFP-VirB T68S. Scale bar represents 1 μm in all images.
